# Salt-free fractionation of complex isomeric mixtures of glycosaminoglycan oligosaccharides compatible with ESI-MS and microarray analysis

**DOI:** 10.1101/759993

**Authors:** Hao Liu, Apoorva Joshi, Pradeep Chopra, Lin Liu, Geert-Jan Boons, Joshua S. Sharp

## Abstract

Heparin and heparan sulfate (Hp/HS) are linear complex glycosaminoglycans which are involved in diverse biological processes. The structural complexity brings difficulties in separation, making the study of structure-function relationships challenging. Here we present a separation method for Hp/HS oligosaccharide fractionation with cross-compatible solvent and conditions, combining size exclusion chromatography (SEC), ion-pair reversed phase chromatography (IPRP), and hydrophilic interaction chromatography (HILIC) as three orthogonal separation methods that do not require desalting or extensive sample handling. With this method, the final eluent is suitable for structure-function relationship studies, including tandem mass spectrometry and microarray printing. Our data indicate that high resolution is achieved on both IPRP and HILIC for Hp/HS isomers. In addition, the fractions co-eluted in IPRP could be further separated by HILIC, with both separation dimensions capable of resolving some isomeric oligosaccharides. We demonstrate this method using both unpurified reaction products from isomeric synthetic hexasaccharides and an octasaccharide fraction from enoxaparin, identifying isomers resolved by this multi-dimensional separation method. We demonstrate both structural analysis by MS, as well as functional analysis by microarray printing and screening using a prototypical Hp/HS binding protein: basic-fibroblast growth factor (FGF2). Collectively, this method provides a strategy for efficient Hp/HS structure-function characterization.

## Introduction

Heparin and heparan sulfate (Hp/HS) are glycosaminoglycans (GAGs), highly anionic unbranched polysaccharides found on the surface of essentially all mammalian cells. Hp/HS consists of a repeating disaccharide structure either β-D-glucuronic or α-L-iduronic acid (GlcA and IdoA, respectively) and α-D-*N*-acetyl-D-glucosamine (GlcNAc), all connected by 1→4 linkages^1^. As a part of biosynthesis, monosaccharides can be differentially *N*- and *O*-sulfated after polymerization. The GlcA can be epimerized to IdoA and can be 2-*O*-sulfated, while the GlcNAc can be deacetylated (usually followed by *N*-sulfation) and/or *O*-sulfated at the 6- and/or 3-position. Since these modifications are incomplete and untemplated, there is enormous structural heterogeneity in Hp/HS chains, including many isomeric structures with widely varying and dynamic compositions^2^. Different cell types will often display different HS structures, and these structures can change as part of the cell’s physiological response^3^. This structural diversity mediates a wide range of protein-GAG interactions of varying specificity and affinity. Hp/HS has been involved in a wide and growing array of physiological and pathophysiological processes, usually mediated through interactions between proteins and a subset of Hp/HS structures^4-6^. While the importance of oligosaccharide structure has been established in many functions of Hp/HS, understanding the structures that underlie any of the various functions is highly challenging. Two technologies that have been applied with considerable success in understanding Hp/HS structure-function relationships are mass spectrometry (MS)^7,8^ and Hp/HS microarrays^9,10^. However, the vast diversity, myriad isomeric structures, and difficulties in separation make the application of these tools towards Hp/HS oligosaccharides very challenging. The vast diversity of potential structures (5.3 million octasaccharides) makes complete coverage by synthetic methods impractical^11^. Therefore, the need for efficient separation technique prior to MS or microarray printing that is readily compatible with these techniques is clear^12,13^.

Low molecular weight heparin (LMWH) is a partially depolymerized product of heparin. In the U.S.A., the most widely used LMWH is enoxaparin sodium, which is generated by β-elimination of heparin^14^. The majority of the depolymerization products have a 4,5-unsaturated dehydro-uronic acid residue (ΔUA) at the non-reducing end, which can be detected at 232 nm by a UV detector^15^. Enoxaparin sodium is a good model for the study of separation and characterization of Hp/HS with high-performance liquid chromatography (HPLC). Normally, size exclusion chromatography (SEC) is the first separation method following partial depolymerization of the Hp/HS polysaccharide^16^. Most methods of Hp/HS depolymerization cleave to the non-reducing end of the uronic acid, resulting in oligosaccharide products that differ by a disaccharide unit in length and can be separated by SEC based on the oligosaccharide length^17^. A common follow-up separation is a charge-based separation, strong anion exchange chromatography (SAX), which has a high resolving power for Hp/HS fractionation^18^. However, the high concentration of sodium chloride (normally 1.0 M) used in SAX makes it incompatible with MS or other forms of chromatography that use organic solvents^19^. Thus the isolated components after SAX must be desalted prior to MS or further separation, causing sample loss compared to the limited sample amount.

Ion-pair reversed phase chromatography (IPRP) is another powerful method for Hp/HS fractionation with the addition of ion-pairing agents, such as pentylamine^20^ or tributylamine^21^. Compared with SAX, IPRP is more easily coupled with other separation methods and MS. Hyphenation of IPRP with SEC and time of flight mass spectrometry have greatly improved separation and provided more complete and important structural information of low molecular weight heparin (LMWH)^22^. But the ion-pair agents will easily induce in-source contamination and cause signal suppression when coupled with MS^13^. Some IPRP agents can also interfere with microarray immobilization schemes^23^. Reverse-phase chromatography without the use of strong ion pairing reagents (RP) has also shown strong separation ability for heavily derivatized chondroitin sulfate isomers and heparan sulfate^24,25^, but the derivatization process leaves the oligosaccharides in non-native conditions not suitable for further functional analysis. Thus a more MS- and microarray-friendly orthogonal separation method for native oligosaccharides is preferred to couple with IPRP.

Hp/HS are very hydrophilic with several polar groups, like sulfates, carboxylates, and hydroxyls. Hydrophilic interaction chromatography (HILIC) has been widely used for the chondroitin sulfate (CS), dermatan sulfate (DS), and Hp/HS separation^26-28^. Since HILIC requires only volatile mobile phase components and a pH range readily compatible with LC-MS analysis, it is particularly useful for sample cleanup prior to MS in order to provide further detailed structural analysis^29^. The orthogonality of IPRP and HILIC separation has been illustrated for *N*-linked glycans, suggesting the potential power of this hyphenation into a multi-dimensional LC scheme for Hp/HS^30^.

The structural variability of Hp/HS is responsible for interactions between Hp/HS and a wide variety of proteins^31^. The fibroblast growth factors (FGFs) are the most thoroughly studied proteins, which bind to the extracellular matrix by interacting with Hp/HS^32,33^. Tetra- and hexasaccharides are sufficient to bind FGF2 with high affinity^34^. The 2-*O*-sulfate iduronate and the *N*-sulfate of glucosamine are critical motifs for FGF2 binding. Carbohydrate microarrays are powerful high-throughput screening platforms to discover Hp/HS-proteins interactions^35^. The use of microarrays of fractionated Hp/HS will substantially improve the understanding of Hp/HS-protein interactions.

Here we present a separation method for heparin/HS oligosaccharide fractionation **(Figure 1)** with solvent and separation conditions that are both readily cross-compatible, as well as a final eluent that is amenable to both electrospray (ESI)-MS and microarray immobilization. This method uses SEC, followed by pentylamine/acetic acid IPRP chromatography, and ending with amide-HILIC. These hyphenated HPLC separations increase resolving power, bridge Hp/HS MS structural analysis and microarray functional studies, and are readily compatible with online or offline ESI-MS. Our data highlights the advantages of SEC-IPRP-HILIC for separation and purification of complex mixtures of Hp/HS, establishing this methodology as a useful tool for efficient and high-resolution Hp/HS fractionation.

**Figure 1.**
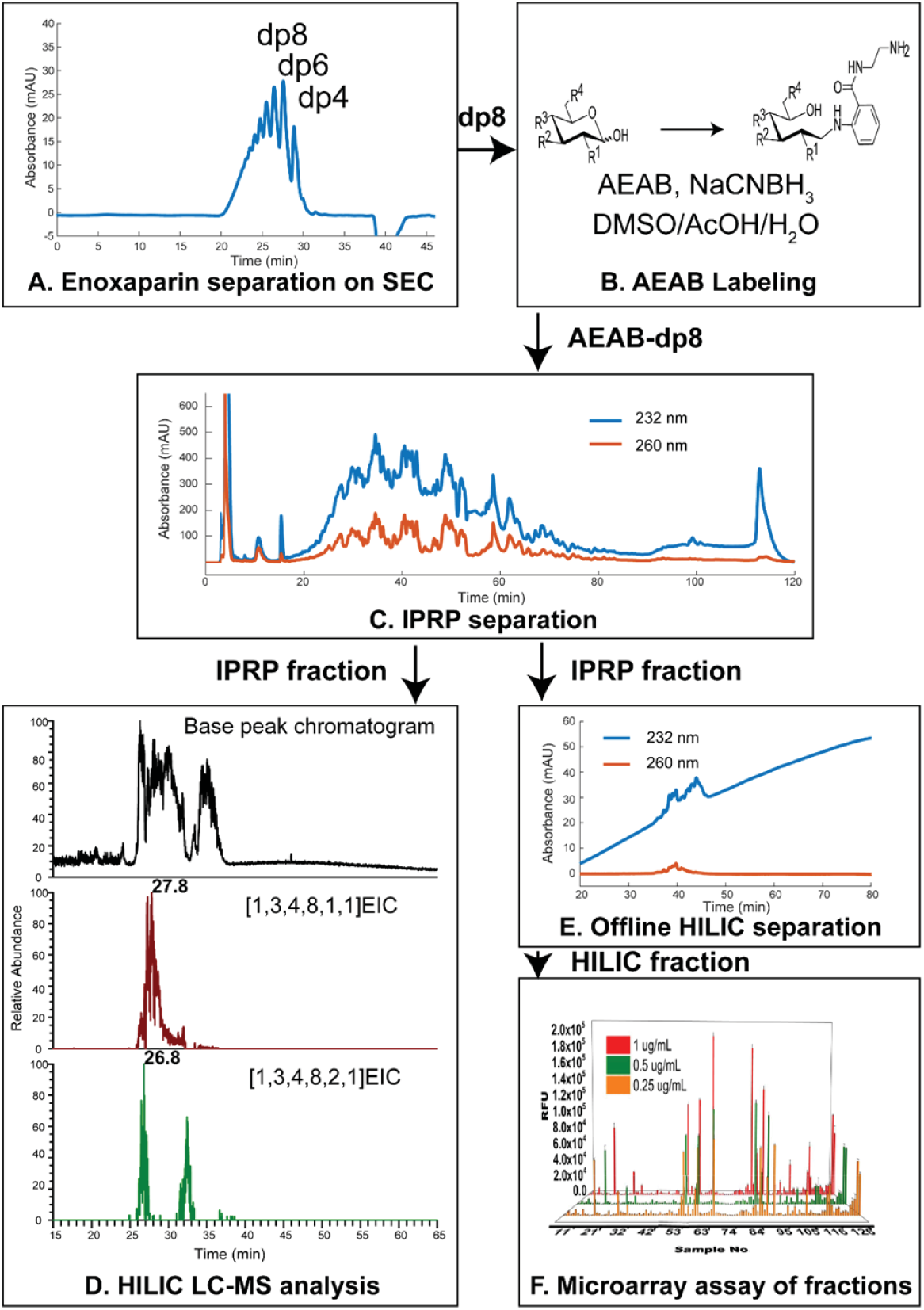
Multi-dimensional separation workflow. Data shown are taken from this study to represent the value of each step. A.) SEC separation of the partially depolymerized Hp/HS, enoxaparin sodium; B.) chemo-selective reductive amination of one SEC fraction (dp8) with AEAB; C.) IPRP separation of AEAB-labeled oligosaccharide SEC fraction (dp8); IPRP fractions could be either analyzed by D.) online HILIC LC/MS (IPRP fraction RT 35 to 36 min shown); or E.) separated by offline HILIC with fractions collected (IPRP fraction RT 36.8 to 38 min shown); F.) microarray immobilization of HILIC fractions and protein binding affinity assay (IPRP and HILIC fraction identities shown in Table S1).

## Results

### The structural complexity of Hp/HS is preserved during AEAB conjugation

To optimize the LMWH sample for both UV detection and microarray immobilization, we used 2-Amino-*N*-(2-aminoethyl) benzamide (AEAB) as a reducing-end tag^36^. In order to ensure we preserve the structural complexity of Hp/HS during AEAB conjugation, we employed the Hp disaccharide standard II-S to develop optimized labeling conditions and validate the chemo-selectivity under the optimized conditions. Hp disaccharide II-S AEAB conjugate was prepared under various amounts of acetic acid and analyzed by HILIC LC-MS. As the amount of acetic acid increased from 10% to 40%, the main product was shifted from conjugation through the aliphatic amine (IIS-AEAB1) to conjugation through the arylamine (IIS-AEAB2, Figure 2). At lower pH, the aliphatic amine was protonated and its nucleophilicity would decrease significantly, leaving the unprotonated arylamine to react with the reducing end of heparin disaccharide II-S to form II-S-AEAB2 conjugate. Therefore, a selective AEAB conjugate is formed with 40% acetic acid, leaving the aliphatic amine free for use in microarray immobilization chemistry. To further confirm the selectivity of the reaction, the synthesized hexasaccharide isomers were labeled under the optimized conditions. The results from HILIC-MS showed that there was only one conjugate LC peak for each isomer under 40% acetic acid (Figure S1). This indicated that the conjugation between the aromatic amine and the reducing end of the sugar was the only detectable product at 40% acetic acid derivatization conditions, excluding the possibilities of introducing isomers during the reductive amination reaction.

**Figure 2.**
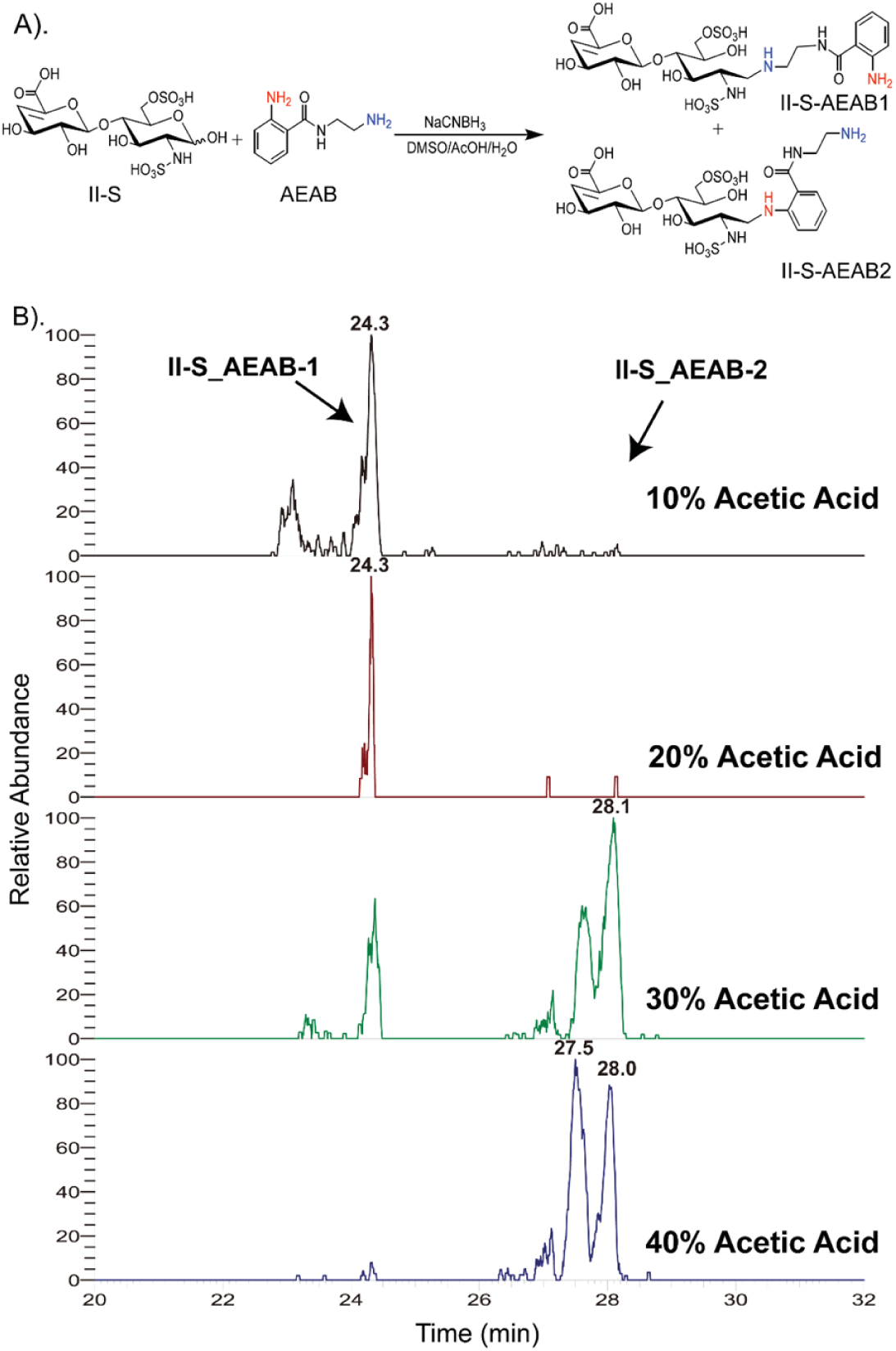
Chemo-selectivity of reductive amination of Hp/HS with AEAB. A.) Scheme of AEAB reductive amination. The difference in pKa between the aromatic amine and the aliphatic amine achieves selective reactivity by controlling the reaction pH. B.) Extracted ion chromatogram of Hp disaccharide II-S: AEAB conjugate after reaction in different concentrations of acetic acid (v/v). The earlier eluting peak represents conjugation of AEAB through the aliphatic amine, with the desired conjugation product through the aromatic amine eluting later.

### Validation of multi-dimensional separation method with synthetic hexasaccharides

We obtained three synthetic hexasaccharides in an unpurified state to evaluate the multi-dimensional separation method, focusing on the separation of isomeric hexasaccharides in a moderately complex background. The expected synthetic structure of U-H-1 is Δ^4,5^UA-GlcNS6S-GlcA-GlcNS6S3S-IdoA2S-GlcNS6S; U-H-2 is Δ^4,5^UA-GlcNS6S-IdoA2S-GlcNS3S-IdoA2S-GlcNS6S; U-H-3 is Δ^4,5^UA-GlcNAc6S-GlcA-GlcNAc6S3S-IdoA2S-GlcNAc6S. Each hexasaccharide was first characterized by amide-HILIC LC-MS and a complex background was detected due to the partial product GAGs in the reaction (Figures S2, S3, and S4). The synthetic hexamers were AEAB-labeled with 40% acetic acid, the optimized condition as described above. These AEAB-labeled synthetic hexasaccharide products were separately injected onto the C18 column with ion-pairing agents so that the retention times (RTs) of three labeled hexaccharides could be confirmed (Figure S5), with the UV trace-based assignments confirmed by HILIC-MS as described below. U-H-3-AEAB (a labeled hexasaccharide with five sulfate groups and three acetyl groups) eluted between 19 min and 20 min. Two isomeric hexasaccharides with eight sulfate groups and no acetyl group, U-H-1-AEAB and U-H-2-AEAB, were separated by IPRP. U-H-2-AEAB eluted between 22 min and 23 min while U-H-1-AEAB eluted between 25 min and 26 min. This overall elution pattern was consistent with IPRP elution order, where lower sulfated oligosaccharides elute earlier. To validate the multi-dimensional separation method, the mixture of three labeled synthetic hexasaccharide products was initially separated by IPRP (Figure 3), and fractions of isomeric hexamers were collected and subsequently separated by online amide-HILIC LC-MS. LC-MS displayed two peaks with an [M −4H]^4-^ ion of m/z 452.5 corresponding to the [1,2,3,8,0,1] (labeled isomeric composition) from two IPRP fractions (RT 22 to 23 min & RT 25 to 26 min) (Figure 4). In the extracted ion chromatograms (EICs), one isomer in IPRP RT 22 to 23 min fraction was eluted at 27.1 min on amide-HILIC, which was U-H-2-AEAB. The other isomer in IPRP RT 25 to 26 min fraction was eluted at 28.0 min on amide-HILIC, which was U-H-1-AEAB. Both isomers were co-eluted on IPRP from 23 to 24 min. The AEAB labeled isomers were not only separated by IPRP (RT 22 to 23 min vs. 25 to 26 min) but LC-MS of the IPRP fraction collected in the valley of the two unresolved isomeric peaks (RT 23 to 24 min) displayed two resolved peaks on HILIC-MS. This result demonstrated the ability of HILIC to separate isomers with distinct sulfate positions under the conditions used here. Overall, IPRP coupled with amide-HILIC not only exchanges solvent to more compatible volatile one for MS sequencing, but also provides high-resolution separations in each dimension for Hp/HS oligosaccharides.

**Figure 3.**
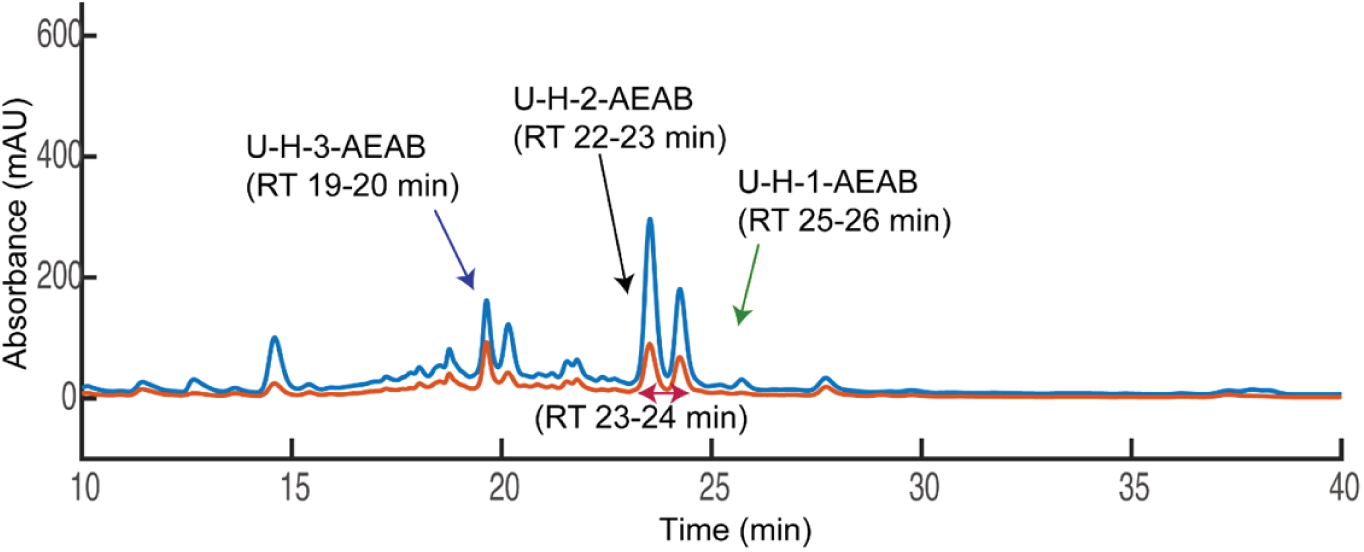
IPRP chromatogram of AEAB labeled synthetic hexasaccharide mixture with UV detection at (blue) 232 nm and (orange) 260 nm. After labeling with AEAB, these synthetic hexasaccharides were separated on IPRP. Peaks were initially assigned based on the elution time of the major peak in the individual synthetic reaction product (Supplementary Information Figure S5). HILIC LC-MS data from fractions shown in black, red, and green are shown in Figure 4.

**Figure 4.**
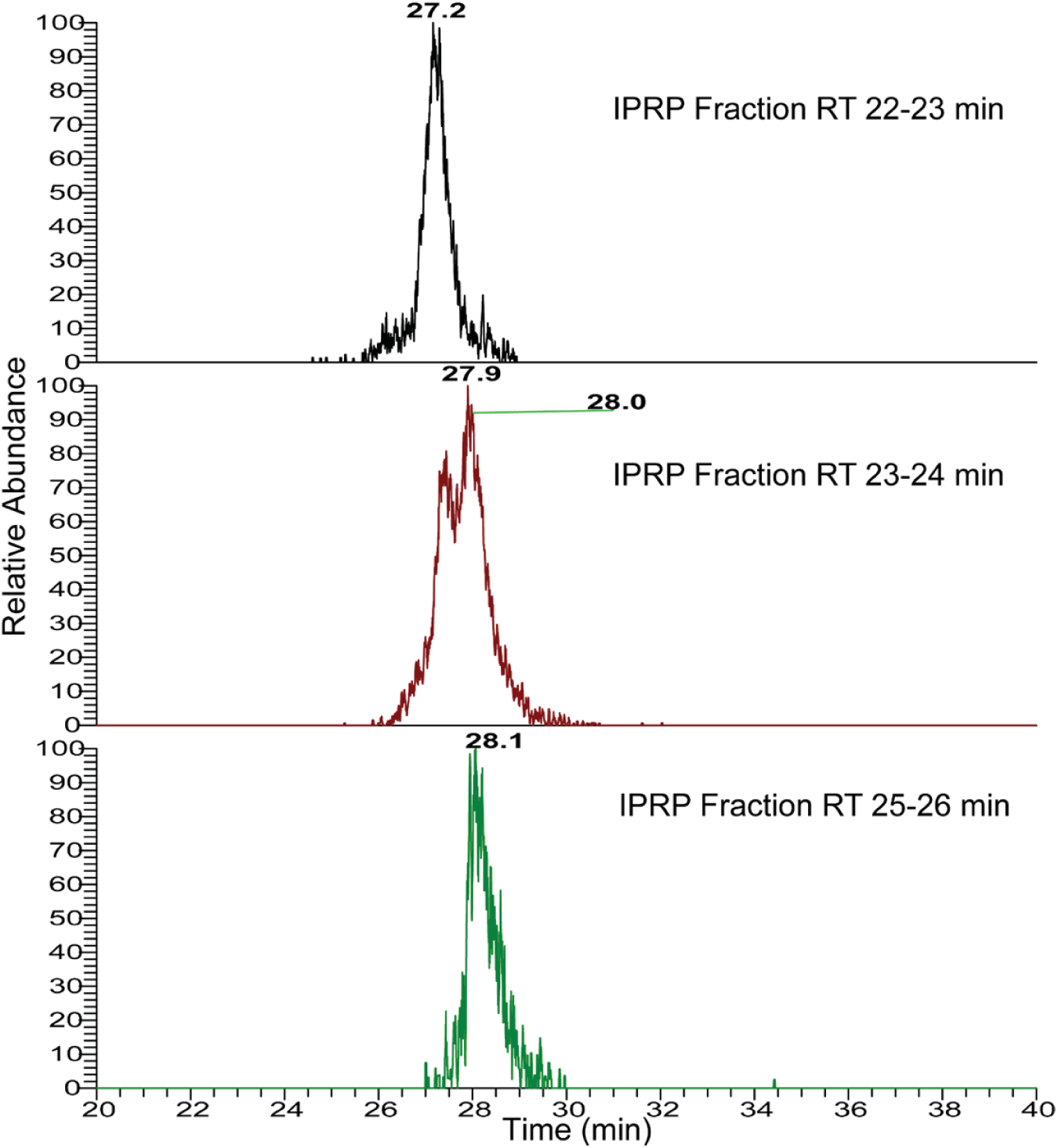
The extracted ion chromatograms of [M −4H]^4-^ ion corresponding to AEAB-labeled [1,2,3,8,0,1] from IPRP fractions. The EICs have shown that the derivatized hexasaccharide composition [1,2,3,8,0,1] collected from IPRP in the 22 to 23 min fraction (top) eluted from the HILIC column around 27.1 min, while the same composition from the IPRP 25 to 26 min fraction (bottom) eluted around 28.0 min on HILIC. The IPRP 23 to 24 min fraction contains the shoulders of both isomer peaks in the IPRP chromatogram, and by HILIC (middle) both the 27.1 and the 28 min peak are partially resolved.

### SEC-IPRP-HILIC separation of enoxaparin sodium

To further evaluate the multi-dimensional separation method, we applied the method to a more complex starting material, enoxaparin sodium. Enoxaparin sodium is a degraded product of heparin; hence it is a good model for highly complex sulfated oligosaccharides derived from natural sources. Due to the multi-dimensional structural complexity of enoxaparin sodium (chain length, sulfate number, and sulfation pattern), size exclusion chromatography was introduced as the first dimension separation to lower the complexity of size. We employed a size-defined octasaccharide fraction to further validate the multi-dimensional separation method.

The octasaccharide SEC fraction of enoxaparin sodium was derivatized with AEAB under optimized conditions and then subjected to IPRP chromatography (Figure 5). Four diverse fractions from the IPRP chromatogram were selected and separately injected onto the amide-HILIC column. The top four possible octasaccharide compositions in each IPRP fraction were analyzed by GlycoWorkbench and listed in Table 1. For the same octasaccharide compositions (e.g. [1,3,4,8,1,1]), the signal was detected in two disparate IPRP fractions (35 to 36 min and 40 to 41 min), which also had different retention times on amide-HILIC (27.84 min vs. 19.69 min) (Figure 6). Interestingly, the huge difference in retention times (ΔRT=5 min on IPRP and ΔRT=8 min on amide-HILIC) indicated that a very high resolution was achieved with the multi-dimensional separation method. Similar isomeric separations could also be found for [1,3,4,7,1,1] and [1,3,4,9,1,1] (Figure S6 through S7). Thus, oligosaccharide composition analysis demonstrated that the powerful multi-dimensional separation method was able to resolve complex isomeric structures, especially sulfate positional isomers.

**Table 1.** Most abundant compositions of AEAB labeled octasaccharides from multi-dimensional separation.

| IPRP fraction (Retention Time) | Oligosaccharide composition<sup>†</sup> |
| --- | --- |
| 27-28 min | [1,3,4,6,1,1] |
|  | [1,3,4,6,2,1] |
|  | [1,3,4,7,1,1] |
|  | [1,3,4,8,0,1] |
| 35-36 min | [1,3,4,7,1,1] |
|  | [1,3,4,8,1,1] |
|  | [1,3,4,8,2,1] |
|  | [1,3,4,9,1,1] |
| 40-41 min | [1,3,4,8,1,1] |
|  | [1,3,4,9,1,1] |
|  | [1,3,4,10,1,1] |
|  | [1,3,4,10,0,1] |
| 58-61 min | [1,3,4,11,0,1] |
|  | [1,3,4,12,0,1] |

<sup>†</sup>Oligosaccharide compositions are given as [ ΔHexA, HexA, GlcN, SO<sub>3</sub>, Ac, AEAB].

**Figure 5.**
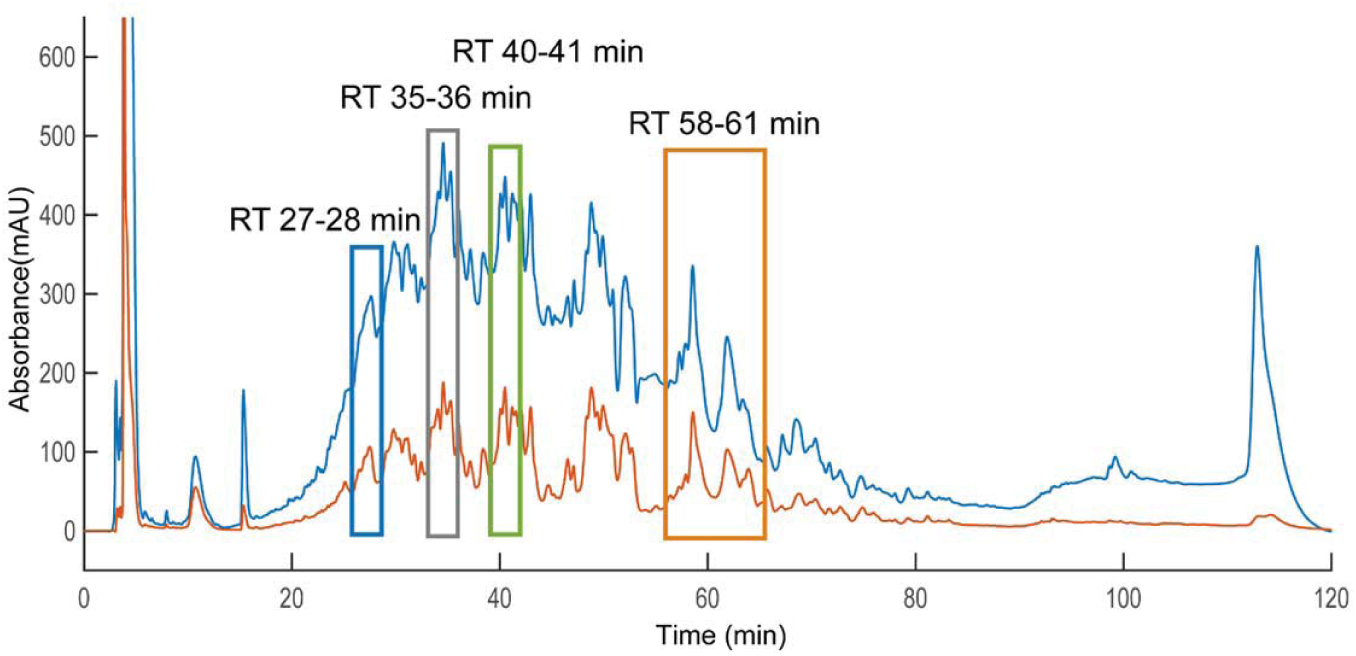
Separation of AEAB labeled enoxaparin octasaccharides using IPRP detected using 232 nm (blue trace) and 260 nm (orange trace) wavelengths. The heterogeneous enoxaparin octasaccharide fraction separated into many poorly-resolved peaks across the elution window. The four fractions highlighted in the chromatogram were collected for further HILIC separation and MS analysis as examples.

**Figure 6.**
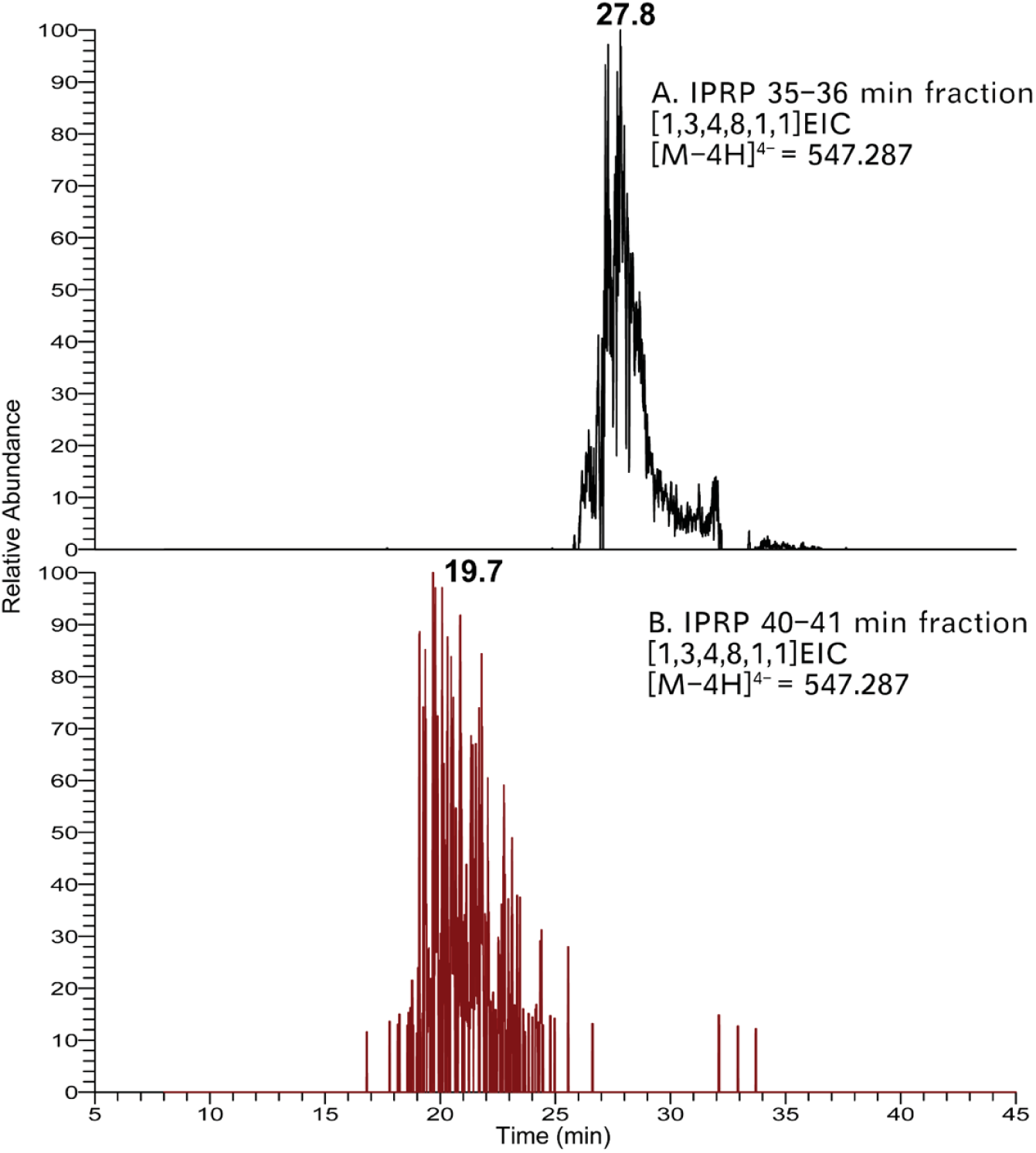
The HILIC EICs of [M −4H]^4-^ ion corresponding to [1,3,4,8,1,1] from IPRP fractions. A.) In the IPRP 35 to 36 min fraction, this composition eluted at 27.8 min. B.) In the IPRP 40-41 min fraction, a different structure with this composition is eluted at 19.7 min. Oligosaccharide compositions are given as [ΔHexA, HexA, GlcN, SO_3_, Ac, AEAB].

Although IPRP itself could separate simple isomers, it may not be sufficient for complex mixtures like Hp/HS. In order to examine the possible advantages of adding in an amide-HILIC dimension, we examined extracted ion chromatograms of various compositions from the surveyed IPRP fractions on HILIC LC-MS as shown in Figures 6 and 7 and Supplementary Information (Figure S8 through S10). For some compositions such as [1,3,4,8,2,1], two isomers co-eluted in IPRP (both in 35-36 min fraction) but were separated on HILIC (26.8 min vs. 32.4 min) (Figure 7). This indicated that the multi-dimensional separation method could provide higher resolution than the one-dimensional separation method for Hp/HS. Additionally, the eluent is completely volatile, making buffer exchange convenient with very little sample loss. Overall, the data validated that the multi-dimensional separation method had a high resolution for Hp/HS, and effectively eliminated ion pairing reagents from the buffer simultaneously.

**Figure 7.**
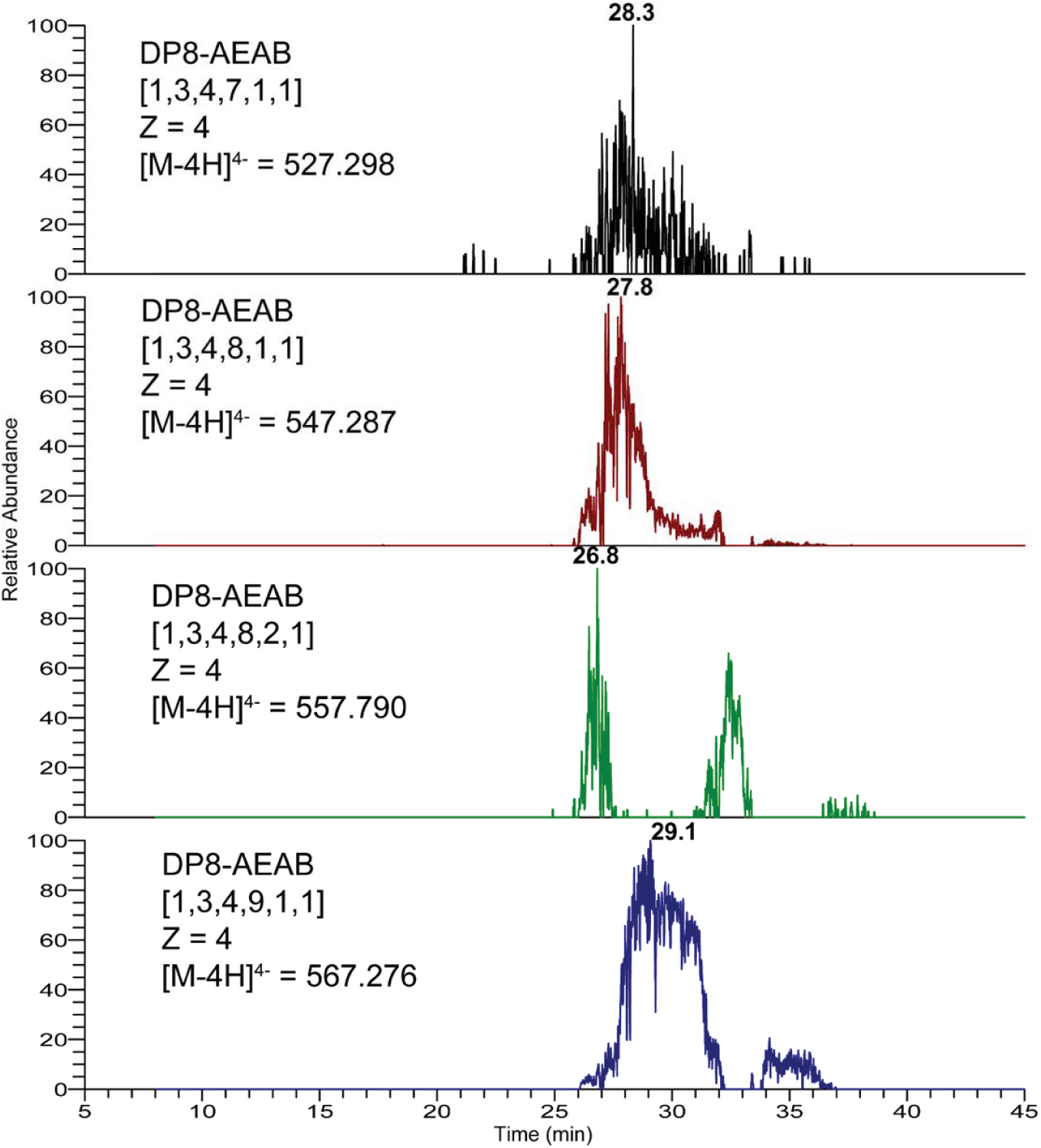
The EICs of the top four octasaccharide compositions from the IPRP 35-36 min fraction. The octasaccharides, which are not separated by IPRP, are resolved by HILIC, including two clearly separated [1,3,4,8,2,1] isomers, illustrating orthogonality between HILIC and IPRP chromatography. Oligosaccharide compositions are given as [ΔHexA, HexA, GlcN, SO_3_, Ac, AEAB].

### The application of the multi-dimensional separation method for microarray functional studies

As a further test of the application of the multi-dimensional separation method, we probed fractions for binding to FGF2. All FGFs have a high affinity for Hp/HS and the affinity of FGF is modulated by different Hp/HS structures^37^. Therefore, this assay would identify which highly enriched fractions had a higher affinity to FGF2 and demonstrate the compatibility of the multi-dimensional separation method with amine-mediated microarray immobilization.

We employed the multi-dimensional separation method described above to lower the structural complexity gradually and constructed Hp/HS oligosaccharide fraction microarrays for functional study, starting with the octasaccharide fraction from SEC. More than 40 fractions were collected from the IPRP run (starting from retention time 24 min to 77 min). HILIC separation was performed for each IPRP fraction, with each peak collected as a fraction and lyophilized. Using AEAB absorbance to quantify oligosaccharide concentration (using a calibration curve based on AEAB-labeled Hp disaccharide), 5nmol of each eluent fraction was aliquoted out and diluted in appropriate buffer to yield a printing concentration of 100 μM^38^. Together with three controls [AEAB labeled enoxaparin sodium octasaccharides (in duplicate), and synthetic octasaccharide], 129 highly enriched oligosaccharide fractions were printed in replicates of 6 on NHS-ester activated glass slides, generating a series of Hp/HS octasaccharide microarrays. Through multi-dimensional separations, some octasaccharide fractions (Sample No. 52, 58, 65, and 85) had a higher affinity to FGF2 with higher fluorescence signal while some octasaccharide fractions (Sample No. 53, 55, 60, and 70) had low affinity to FGF2 with lower fluorescence signal (Figure 8, Figure S11 and Table S1). Analysis of FGF2 binding versus retention time by IPRP and HILIC reveals a strong correlation between HILIC retention and FGF2 binding (Figure 9). Previous work by Zaia and coworkers have shown that amide-HILIC retention time is dominated by acetyl and sulfate composition for Hp/HS oligosaccharides of a given size ^39^, indicating that FGF2 binding increases as sulfate content increases. This is consistent with previous microarray results using synthetic oligosaccharides that showed that FGF2 binds all highly sulfated GAG tetrasaccharides well, while intermediately sulfated structures showed a clear structure-function relationship ^40^, and is consistent with other FGF2 binding structure-function studies showing preference for high sulfation and indicating the importance of 2-*O*-sulfation and *N*-sulfation to FGF2 binding ^41-44^.

**Figure 8.**
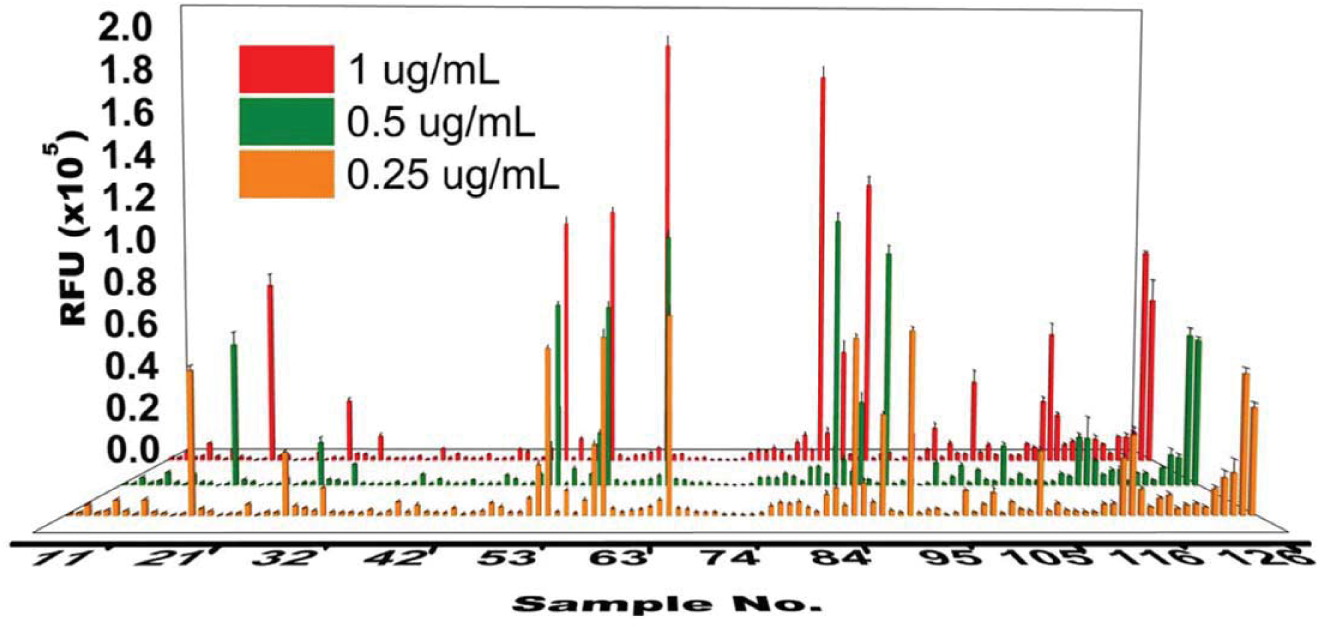
FGF2-binding assay on Hp/HS octasaccharide-AEAB microarray Each fraction is printed in replicates of 6 in a single block with droplet volume around 400 pL. Fractions from the multi-dimensional separation method are directly applicable to microarray study. Based on the fluorescent intensity of each spot, it indicates that each fraction has different binding affinities to FGF2. Fractions from No. 1 to No. 126 were eluents from the multi-dimensional separation method. Fractions No. 127 & 128 were unfractionated octasaccharide. Fraction No. 129 was synthetic octasaccharide as a positive control. An image of the microarray is in Supplementary Information Figure S11. Detailed information of each fraction is listed in Table S1.

**Figure 9.**
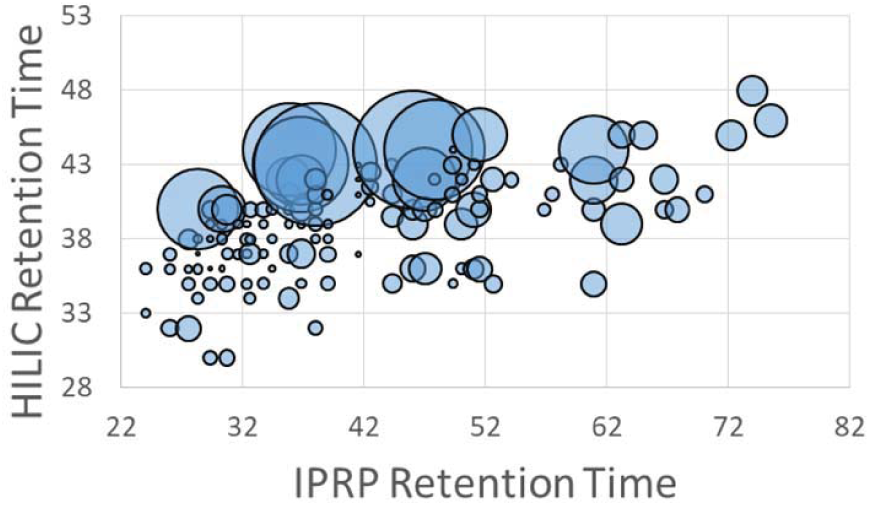
Correlation between IPRP retention time, HILIC retention time, and FGF2 binding response by microarray analysis at 1 μg/mL, represented by bubble size. A strong correlation between HILIC retention time and FGF2 binding is apparent, indicating that FGF2 is binding ligands that are more highly sulfated. No correlation between IPRP retention time and FGF2 binding is apparent.

## Discussion

Although Hp/HS influence numerous physiological and pathophysiological processes, their high degree of size polydispersity and sequence microheterogeneity make the structure-function characterization challenging. In most modern cases, structural analysis is limited to compositional analysis, with oligosaccharide sequences rarely addressed from complex mixtures. Here we demonstrate a three-dimensional LC separation method, SEC-IPRP-HILIC, that is applied for the efficient analysis of Hp/HS with high resolution separation of isomeric sequences, compatibility with both on-line and off-line MS structural analysis and microarray functional analysis.

We have shown that IPRP and HILIC offer orthogonal selectivity with a resolution that can, at least in some cases, separate positional isomers that differ by the site of modification and possibly uronic acid stereochemistry (which was not directly probed in this study), with an overview of the contribution of each step demonstrated in **Figure 1**. Although IPRP itself could separate Hp/HS isomers, our data demonstrate incomplete separation of some isomers, indicating that extra separation is needed to resolve more detailed structural information in Hp/HS. For the first time, clean HILIC separation of isomeric Hp oligosaccharides is demonstrated, as shown in Figure 6 for composition [1,3,4,8,1,1] and octasaccharide compositions [1,3,4,7,1,1] and [1,3,4,9,1,1] in Supplementary Information (Figure S6 and S7) among others. Each separation dimension adds resolution to the overall separation, in addition to HILIC serving as a desalting step to remove ion pairing reagents from IPRP fractions.

We also demonstrate the compatibility of the multi-dimensional separation method to glycan microarray analysis. The fractions from HILIC are directly compatible with microarray immobilization. Ammonium formate, acetonitrile and water used in our HILIC method are volatile and does not interfere with the immobilization method and allows buffer exchange by evaporation or lyophilization, with very little sample loss. Glycan microarrays permit the study of specific Hp/HS-binding proteins quantitatively even with a very limited amount. AEAB incorporates stoichiometrically to oligosaccharides by specific reductive amination to provide both increased UV detection sensitivity and a reactive group for immobilization, and only 1 to 2 nmols of AEAB-labeled Hp/HS fractions are required to prepare microarrays. Our unfractionated octasaccharide microarray results (Sample No. 127 & No. 128) showed lower binding than our fractions with highest activity, confirming that multi-dimensional separation does not compromise the binding capability of the oligosaccharides. As anticipated, the microarray data also display clear diversity in binding affinities of each fraction (Figure S11 and Table S1), consistent with the intermediate structural specificity of FGF2 binding to Hp/HS oligosaccharides^45-47^. Although Hp/HS separated by the multi-dimensional separation method might have similar compositions, both their separation and diverse binding affinities indicate that they are structurally and functionally different. It is possible that the unique structures or composition motifs binding with FGF2 are separated from unbinding Hp/HS fractions. While AEAB-labeling and microarray analysis provide powerful options for structure-function analysis, this method will also work for salt-free fractionation of depolymerized oligosaccharides without derivatization of the reducing end, so long as the unsaturation at the non-reducing end is present to provide a chromophore or some other LC detection method is available.

Overall, this multi-dimensional separation method provides high-resolution separation of Hp/HS and thus generate Hp/HS microarrays for functional study with high purity. The ability to perform HILIC-MS directly allows investigators to directly couple structural interrogations with functional analysis. Future studies will couple this separation method with more detailed Hp/HS structural interrogation methods that determine precise sites of sulfation ^48-50^, which are usually lost in MS/MS analysis. In addition, such Hp/HS microarrays will be useful for screening more Hp/HS-binding proteins and will permit the discovery of new protein-carbohydrate interactions. Thus, this method is invaluable in recognizing complex protein-carbohydrate interactions and is also essential to reveal functional Hp/HS structures as novel biomaterials or therapeutics.

## Methods

### Chemicals and reagents

Enoxaparin sodium injection (Winthrop/Sanofi, Bridgewater, NJ, USA) was used as provided. 2-amino-*N*-(2-aminoethyl)-benzamide (AEAB) was purchased from Chem-Impex International (Wood Dale, IL, USA). Water was purified by a MilliQ^®^ system (Millipore, Bedford, MA, USA). Ammonium formate and sodium cyanoborohydride were purchased from Thermo Fisher Scientific (Waltham, MA, USA). Glacial acetic acid, dimethyl sulfoxide (DMSO), formic acid, and pentylamine were purchased from Millipore Sigma (St. Louis, MO, USA). All chemicals used were at least of analytical grade. Three hexasaccharides with unsaturated uronic acid at the non-reducing end were chemically synthesized using modified reported procedures and the reaction products were desalted over a Bio-Gel P-2 column (Bio-Rad, Hercules, CA, USA) without further purification; the detailed synthetic methodology will be reported elsewhere. The expected synthetic structure of U-H-1 is Δ^4,5^UA-GlcNS6S-GlcA-GlcNS6S3S-IdoA2S-GlcNS6S; U-H-2 is Δ^4,5^UA-GlcNS6S-IdoA2S-GlcNS3S-IdoA2S-GlcNS6S; U-H-3 is Δ^4,5^UA-GlcNAc6S-GlcA-GlcNAc6S3S-IdoA2S-GlcNAc6S. Recombinant human fibroblast growth factor basic was purchased from R&D Systems, Inc. (Minneapolis, MN, USA). Rabbit anti-FGF-2 antibody was purchased from Abcam (Cambridge, MA, USA). Goat anti-rabbit Cy5 antibody was purchased from Life Technologies (Carlsbad, CA, USA).

### Size exclusion chromatography (SEC)

SEC separation was performed on a chromatographic column packed with P-10 media (Bio-Rad^®^, 45-90 μm) and collected by a Frac-920 fraction collector (GE Healthcare Life Sciences, Chicago, IL, USA). 100 mg of commercial Enoxaparin Sodium Injection was injected onto the SEC column and separated by using 0.5 M ammonium bicarbonate eluent at a flow rate of 0.2 mL/min. GAG oligosaccharide quantities were measured using a NanoDrop™ 2000 spectrophotometer (Thermo Fischer Scientific) at a detection wavelength of 232 nm. Fractions from each peak were pooled and evaporated with dry nitrogen gas flow.

### Preparation of AEAB oligosaccharide derivatives

The enoxaparin sodium size-fractionated samples were re-dissolved with water and acetic acid. 0.4 M AEAB in DMSO and 1 M sodium cyanoborohydride in DMSO were added. The ratio of DMSO and acetic acid was varied with acetic acid contents of 10%, 20%, 30% and 40% v/v tested, with 40% acetic acid used as an optimized condition. The mixture was then incubated at 37 °C overnight and dried with nitrogen gas. The dried sample was resuspended in water/methanol (80:20, v/v). It was analyzed and purified by HPLC SEC column (4.6 mm × 300 mm, 1.7 μm, Waters) using a Dionex UltiMate 3000 LC system (Thermo Fisher Scientific) following the method of Dr. Zaia and coworkers^51^. Based on high-resolution SEC separation of derivatized II-S disaccharide standard, the reductive amination reaction had a yield of ∼99% at 40% acetic acid (data not shown). Fractions were collected with an online auto-sampler and further dried down with nitrogen gas. Each synthetic hexasaccharide followed the same protocol.

### Ion-pair reverse phase chromatography (IPRP)

IPRP separations were performed on a Dionex UltiMate 3000 LC system (Thermo Fisher Scientific) using a C18 column (150 mm × 2.1 mm, 2.6 μm, Kinetex^®^, Phenomenex), with a UV-visible spectrophotometric detector. Phase A was 95% water and 5% acetonitrile and phase B was acetonitrile. The ion-pairing agents were 20 mM pentylamine (PTA) and 20 mM acetic acid. All chromatographic separations were monitored at both 232 nm and 260 nm wavelengths. The separation gradient started with 10% B for 5 min, ramped to 20% B and held for 10 min, followed a linear gradient from 20% to 29% B over 65 min, changed to 50% B over 5 min, washed with 50% B for 25 min, then changed to 10 % B over 1 min and re-equilibrated for 24 min. The flow rate was 0.1 mL/min. The column temperature was 40 °C. Fractions were collected with an online auto-sampler and further dried under the nitrogen gas flow.

### Amide hydrophilic interaction chromatography (HILIC)

Hydrophilic interaction chromatography (HILIC) was performed on a BEH Amide HILIC column (50 mm × 1 mm, 1.7 μm, ACQUITY UPLC^®^ BEH Amide, Waters). The flow rate was set to 0.1 mL/min. The gradient for hexasaccharide analysis started with 90% B for 5 min, with a linear gradient down to 30% B over 60 min, held at 30% B for 4 min and finally washed with 10% B for 10 min and re-equilibrated with 90% B for 10 min. The gradient for octasaccharide analysis started at 75% B for 5 min, with a linear gradient from 75% B to 30% B over 60 min, held at 30% B for 4 min and finally washed with 10% B for 10 min and re-equilibrated with 75% B for 10 min. Buffer A was 10 mM ammonium formate (pH 4.4, adjusted with formic acid) and buffer B was 98% acetonitrile with 2% buffer A for online LC-MS analysis. Buffer A was 50 mM ammonium formate (pH 4.4, adjusted with formic acid) for microarray fractionation.

### Online LC-MS method and MS data analysis

The fractionated samples after IPRP were analyzed by LC-MS on a Thermo Orbitrap Fusion Tribrid (Thermo Fisher Scientific) coupled with a Dionex UltiMate 3000 liquid chromatography (Thermo Fisher Scientific) using a BEH Amide HILIC column (50 mm × 1 mm, 1.7 μm, ACQUITY UPLC^®^ BEH Amide, Waters). The oligosaccharides were analyzed in negative ion mode with the ESI voltage set to 1.8 kV. Full mass scan range was set to 230-1000 m/z at a resolution of 60000, and the automatic gain control (AGC) target was set to 1 × 10^5^ counts. The datasets generated during and/or analysed during the current study are available from the corresponding author on reasonable request. GlycoWorkbench software was used to interpret the possible compositions of Hp/HS oligosaccharide ion signals. To interpret the MS data with GlycoWorkbench, we set searching constraints based on the biosynthetic rules of heparin. The maximum value of dehydro-hexuronic acid was set to 1 while the maximum value for hexuronic acid and hexosamine were both set to 5. The maximum number of sulfate groups was 24 and the maximum number of acetate groups was 5. The reducing end delta mass was selected as a mass value of 163.1 Da for the AEAB label. Compositions are reported according to a modified version of the bracketed nomenclature of Dr. Zaia and coworkers [A, B, C, D, E, F], where A is the number of dehydro-uronic acid monosaccharides, B is the number of uronic acid monosaccharides, C is the number of glucosamine monosaccharides, D is the number of sulfations, E is the number of acetylations, and F is the number of AEAB labels in the identified composition^51^.

### Microarray printing, binding assay, and scanning

The arrays were constructed using the guidelines v 1.0. provided by the Minimum Information for a Glycomics Experiment Project (MIRAGE) as reported here^38^.

126 fractions after off-line HILIC separation were printed on NHS-ester activated glass slides (NEXTERION^®^ Slide H, Schott Inc.) using a Scienion sciFLEXARRAYER S3 non-contact microarray equipped with a Scienion PDC80 nozzle (Scienion Inc.). Individual samples (5 nmols) were dissolved in sodium phosphate buffer (50 μL, 0.225 M, pH 8.5) at a concentration of 100μM and were printed in replicates of 6 with spot volume ∼ 400 pL, at 20°C and 50% humidity. Each slide has 24 subarrays in a 3×8 layout. After printing, slides were incubated in a humidity chamber for 8 hours and then blocked for one hour with a 5 mM ethanolamine in a Tris buffer (pH 9.0, 50 mM) at 50°C. Blocked slides were rinsed with DI water, spun dry, and kept in a desiccator at room temperature for future use.

Printed glass slides were incubated with premixed solution of recombinant human fibroblast growth factor basic (1.0 to 0.25 μg/mL FGF-2), rabbit anti-FGF-2 antibody (1:300) and goat anti-rabbit Cy5 antibody (1:300) at room temperature in the dark. After 1h, the slide was sequentially washed by dipping in TSM wash buffer (2 min, containing 0.05 % Tween 20), TSM buffer (2 min) and, water (2 × 2 min), spun dry. The slides were scanned using a GenePix 4000B microarray scanner (Molecular Devices) at the appropriate excitation wavelength with a resolution of 5 μm. Various gains and PMT values were employed in the scanning to ensure that all the signals were within the linear range of the scanner’s detector and there was no saturation of signals. The image was analyzed using GenePix Pro 7 software (version 7.2.29.2, Molecular Devices).

The data was analyzed with our home written Excel macro to provide the results. The highest and the lowest value of the total fluorescence intensity of the six replicates spots were removed, and the four values in the middle were used to provide the mean value and standard deviation.

## Acknowledgments

This research is supported by the National Institute of General Medical Sciences as part of the Research Resource for integrated Glycotechnology (P41GM103390). We thank Waters for donating the BEH Amide HILIC column used in this study.

## Author contributions statement

H.L. performed all fractionation experiments and interpreted all LC and MS data. J.S.S. developed this experimental model and analyzed LC and MS data. A.J., P.C. and G-J.B. synthesized three hexasaccharides. L.L., P.C. and G-J.B. performed the FGF2 binding affinity assay with microarrays. All authors participated in the writing of this manuscript.

## Data Availability

The data generated and analyzed during the current study are available on request.

