## Supplementary Information for "Salt-free fractionation of complex isomeric mixtures of glycosaminoglycan oligosaccharides compatible with ESI-MS and microarray analysis"

Hao Liu[1], Apoorva Joshi[2,3], Pradeep Chopra[3], Lin Liu[3], Geert-Jan Boons[2,3,4], Joshua S. Sharp[1],

1. Department of BioMolecular Sciences, University of Mississippi, Oxford, MS 38677, USA.

2. Department of Chemistry, University of Georgia, Athens, GA 30602, USA.

3. Complex Carbohydrate Research Center, University of Georgia, Athens, GA 30602, USA.

4. Department of Chemical Biology and Drug Discovery, Utrecht Institute for Pharmaceutical Sciences, and Biivoet Center for Biomolecular Research, Utrecht University, Universiteitsweg 99, 3584 CG Utrecht, The Netherlands.

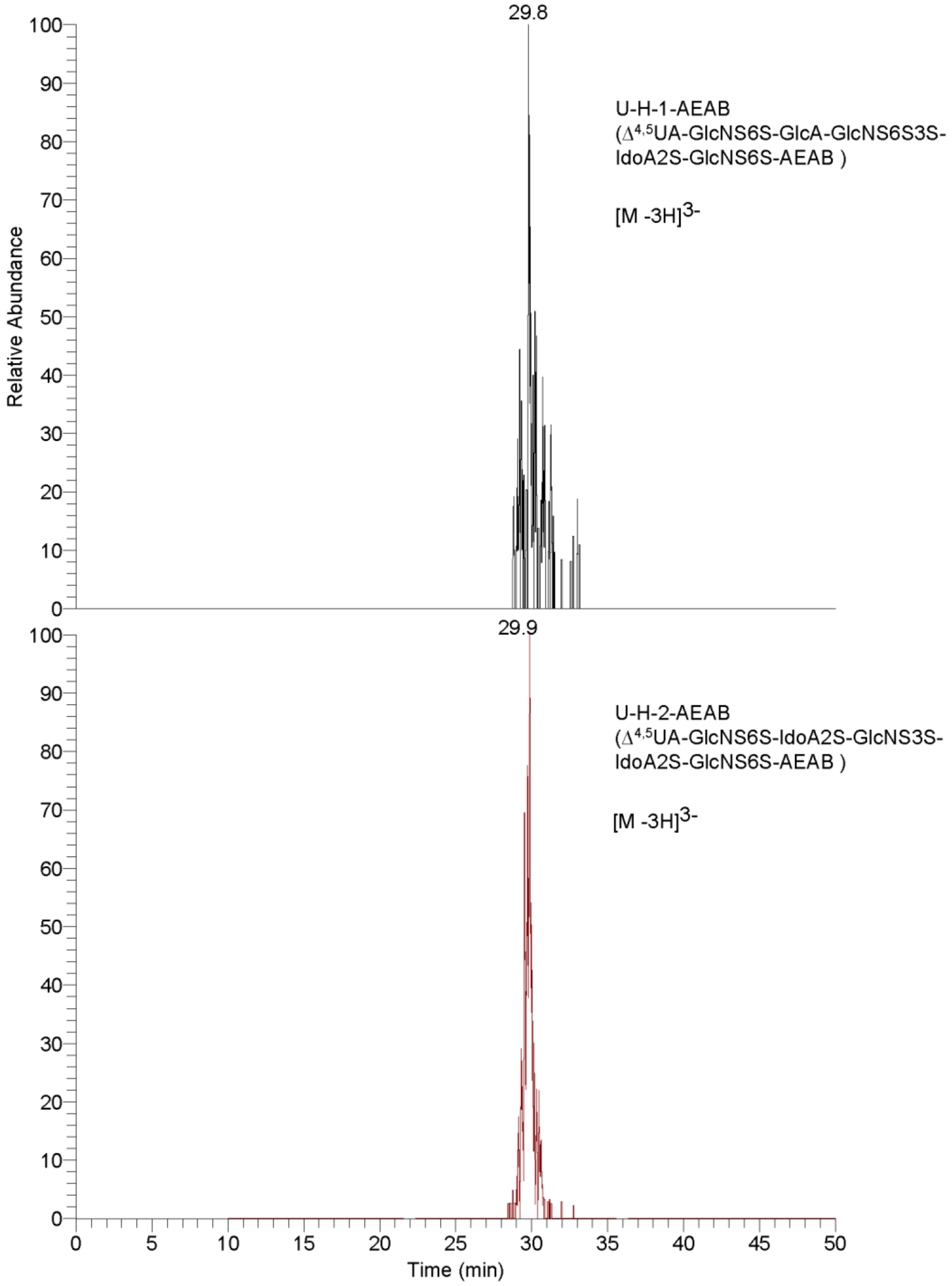


**Figure S1.**

Amide-HILIC LC-MS extracted ion chromatogram (EIC) data for each AEAB labeled synthetic hexasaccharide isomer. With optimized conditions, there is only one labeled product for each synthetic hexasaccharide.

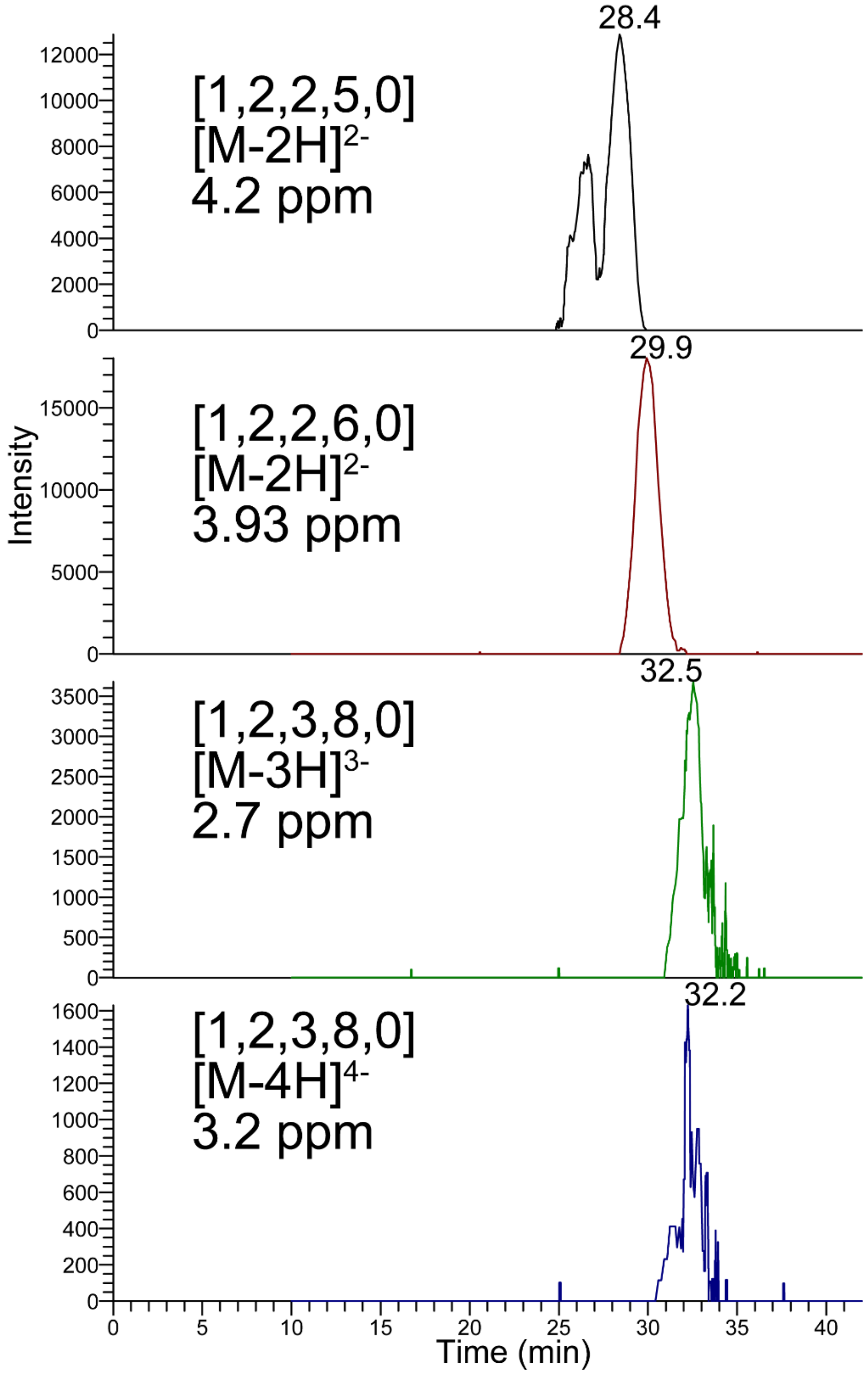


**Figure S2.**

Impurities detected in the HILIC separation of synthetic hexasaccharide, U-H-1, [1, 2, 3, 8, 0]. Oligosaccharide compositions are given as *[ΔHexA, HexA, GlcN,* $SO_3$*, Ac].*

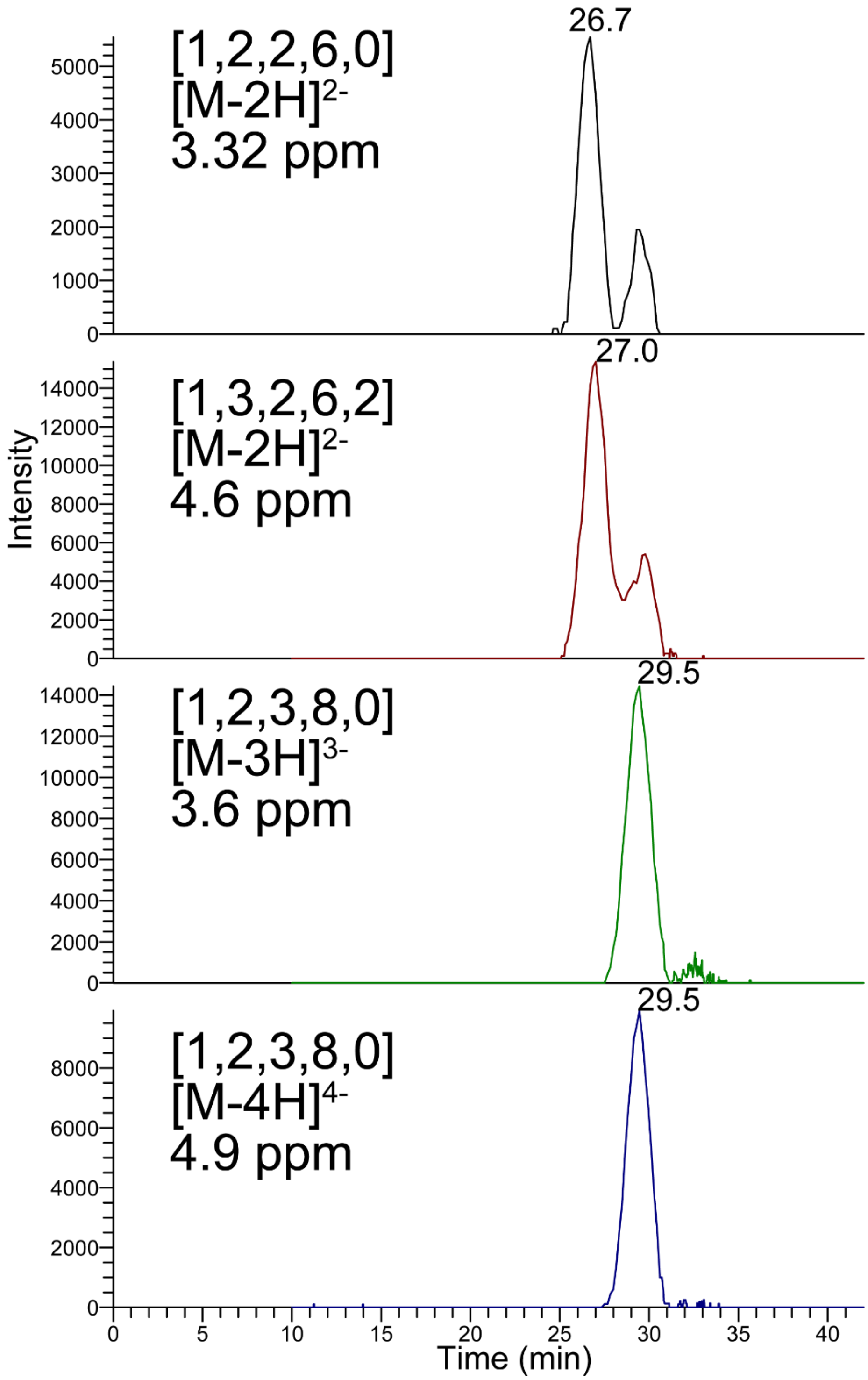


**Figure S3.**

Impurities detected in the HILIC separation of synthetic hexasaccharide, U-H-2, [1, 2, 3, 8, 0].

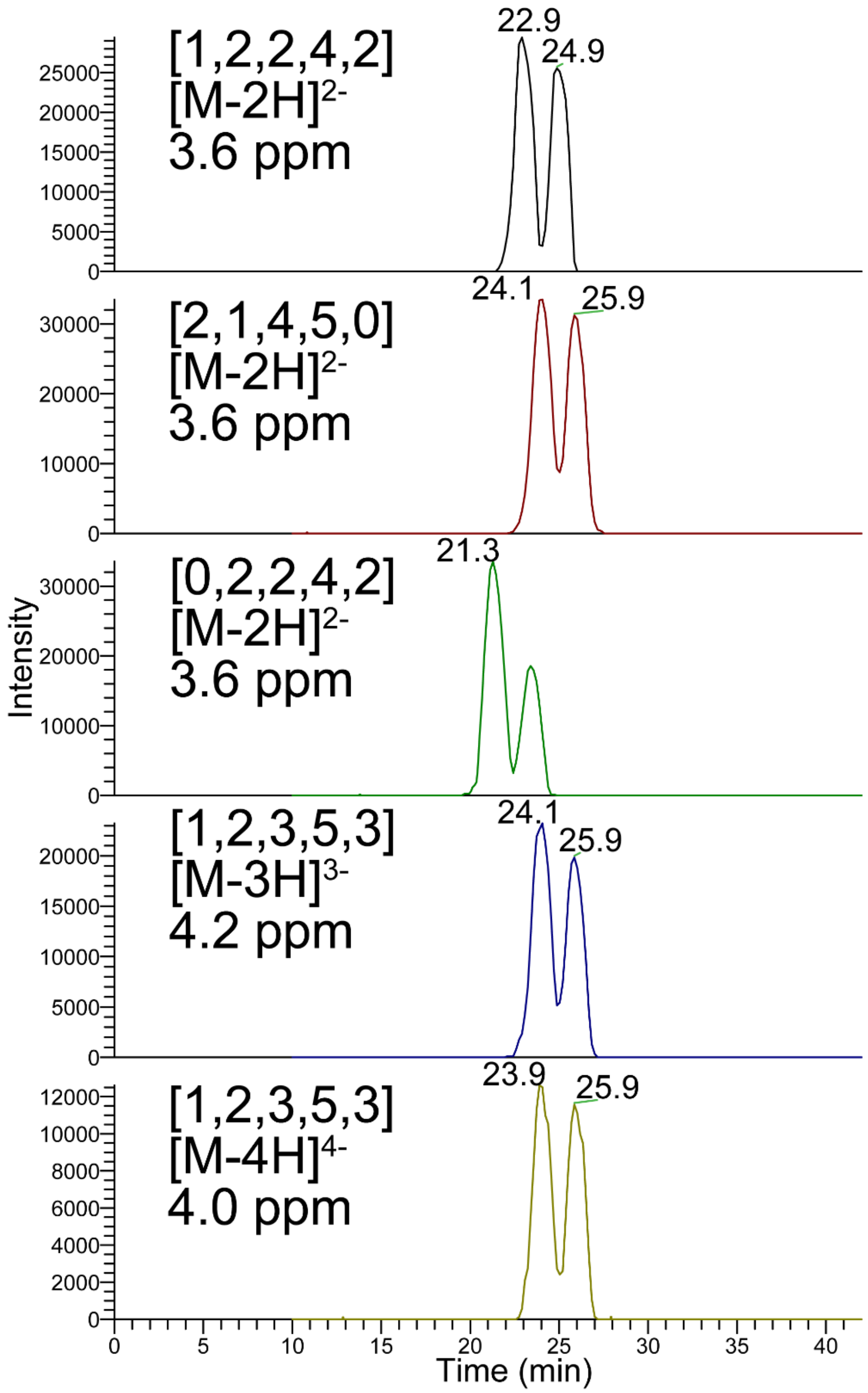


**Figure S4.**

Impurities detected in the HILIC separation of synthetic hexasaccharide, U-H-3, [1, 2, 3, 5, 3].

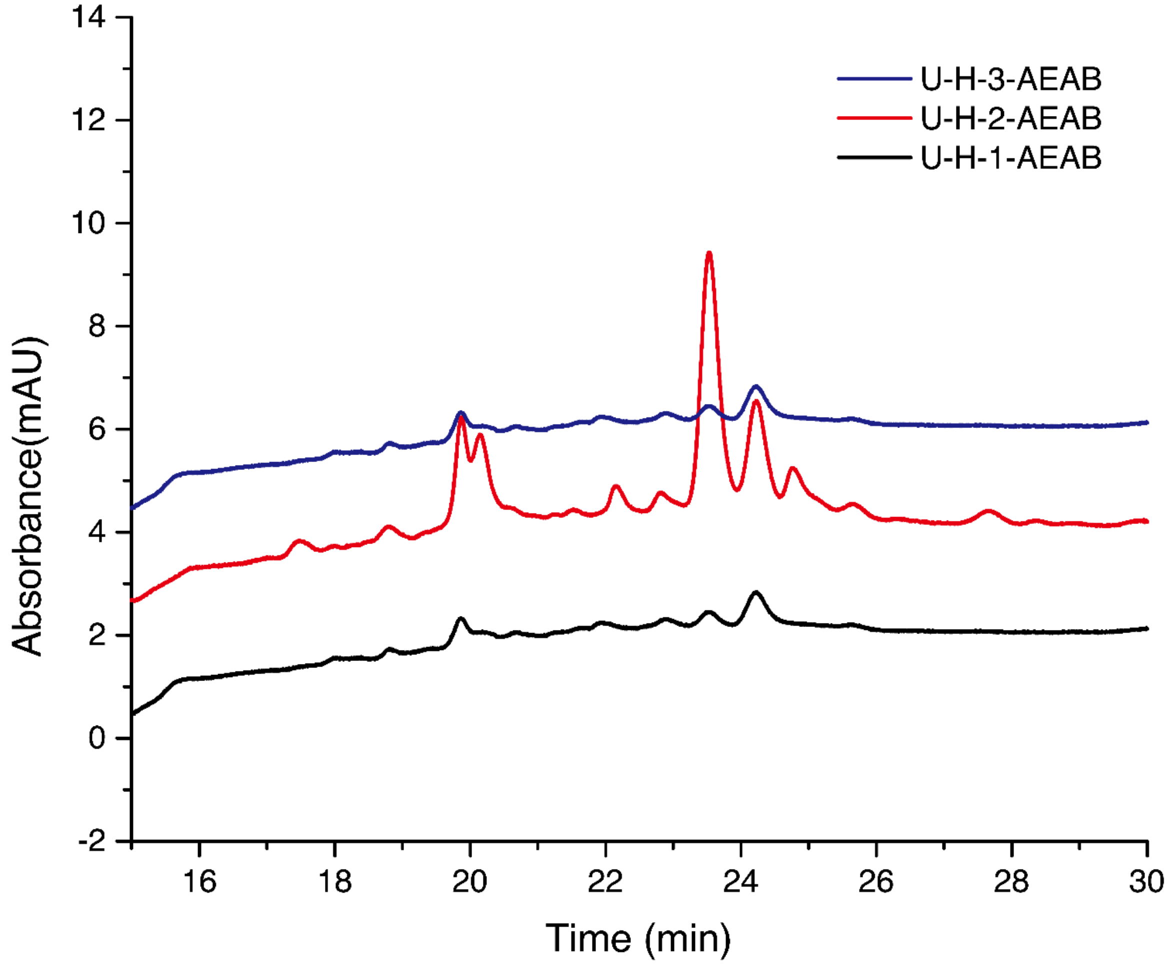


**Figure S5.**

IPRP Analysis of AEAB-labeled unpurified synthetic hexasaccharides injected separately. The U-H-1-AEAB was mainly eluted after 24 min while U-H-2-AEAB was mainly eluted between 22 to 23 min. Non-isomeric *N*-acetylated hexamer U-H-3-AEAB eluted earlier than either isomer.

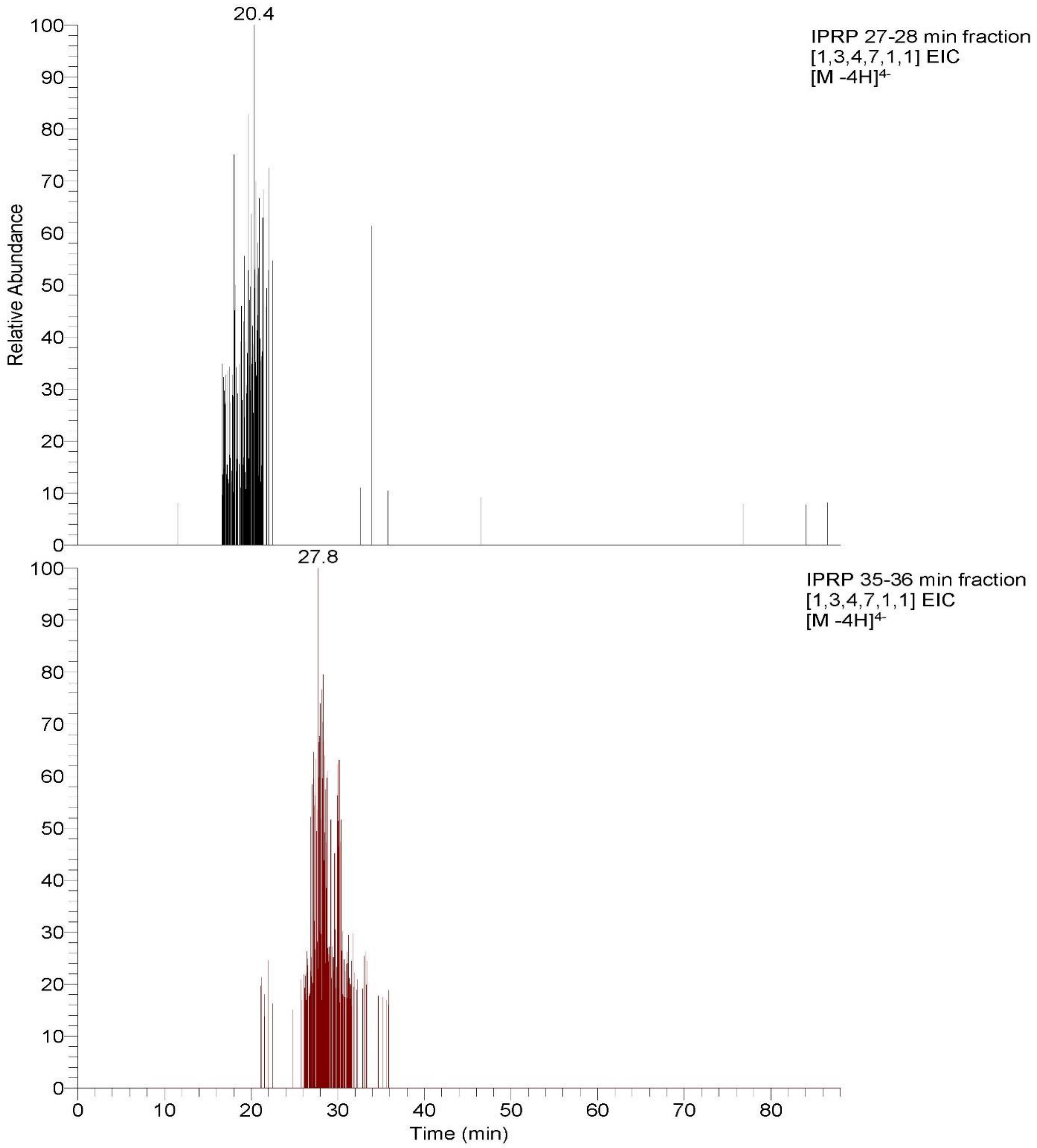


**Figure S6.**

The EICs of [1,3,4,7,1,1] from enoxaparin dp8 (top) IPRP 27-28 min fraction and (bottom) 35-36 min fraction.

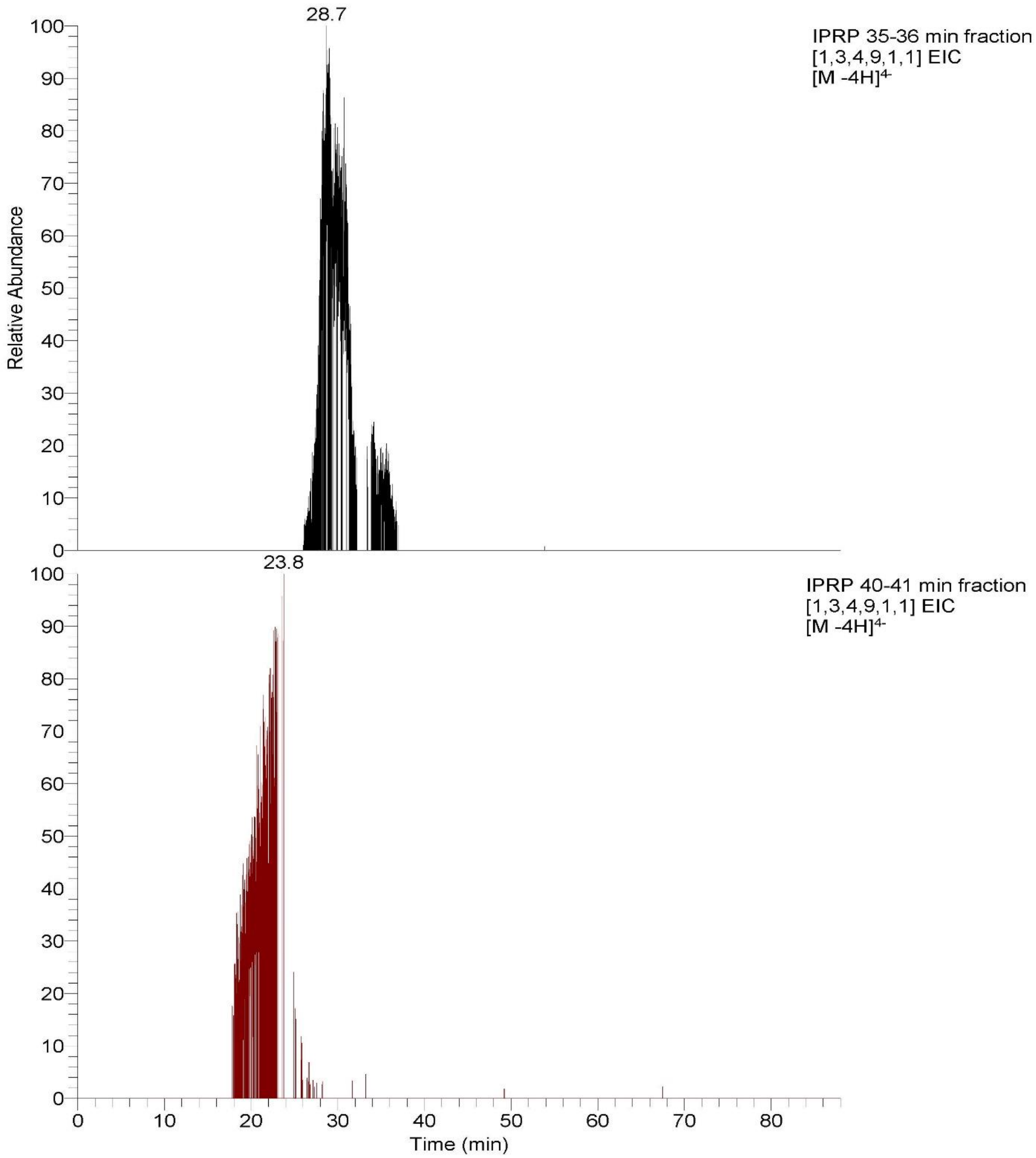


**Figure S7.**

The EICs of [1,3,4,9,1,1] from enoxaparin dp8 (top) IPRP 35-36 min fraction and (bottom) 40-41 min fraction.

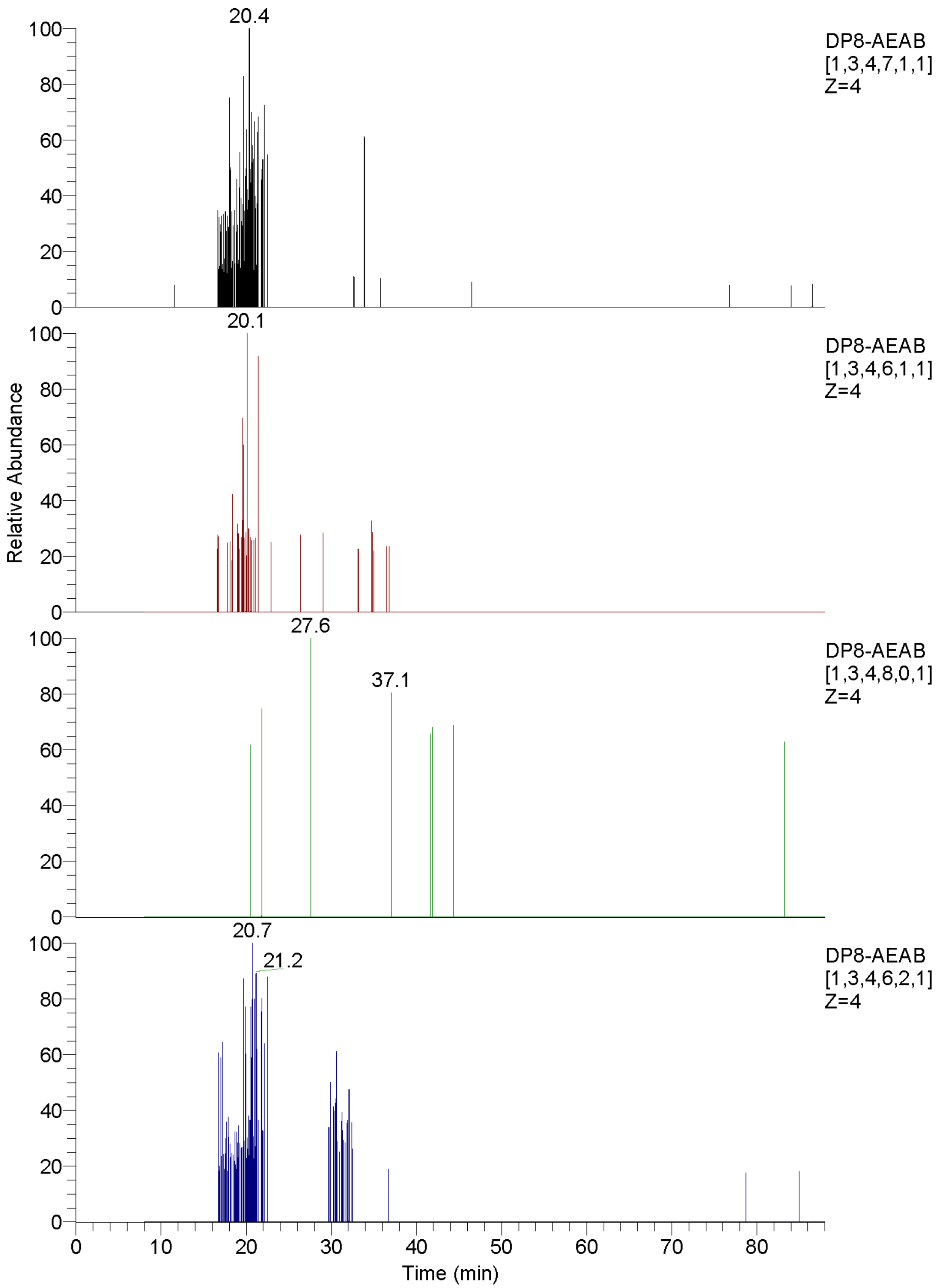


**Figure S8.**

The HILIC-MS EICs of the top four possible octasaccharide compositions of enoxaparin dp8 from the IPRP 27-28 min fraction.

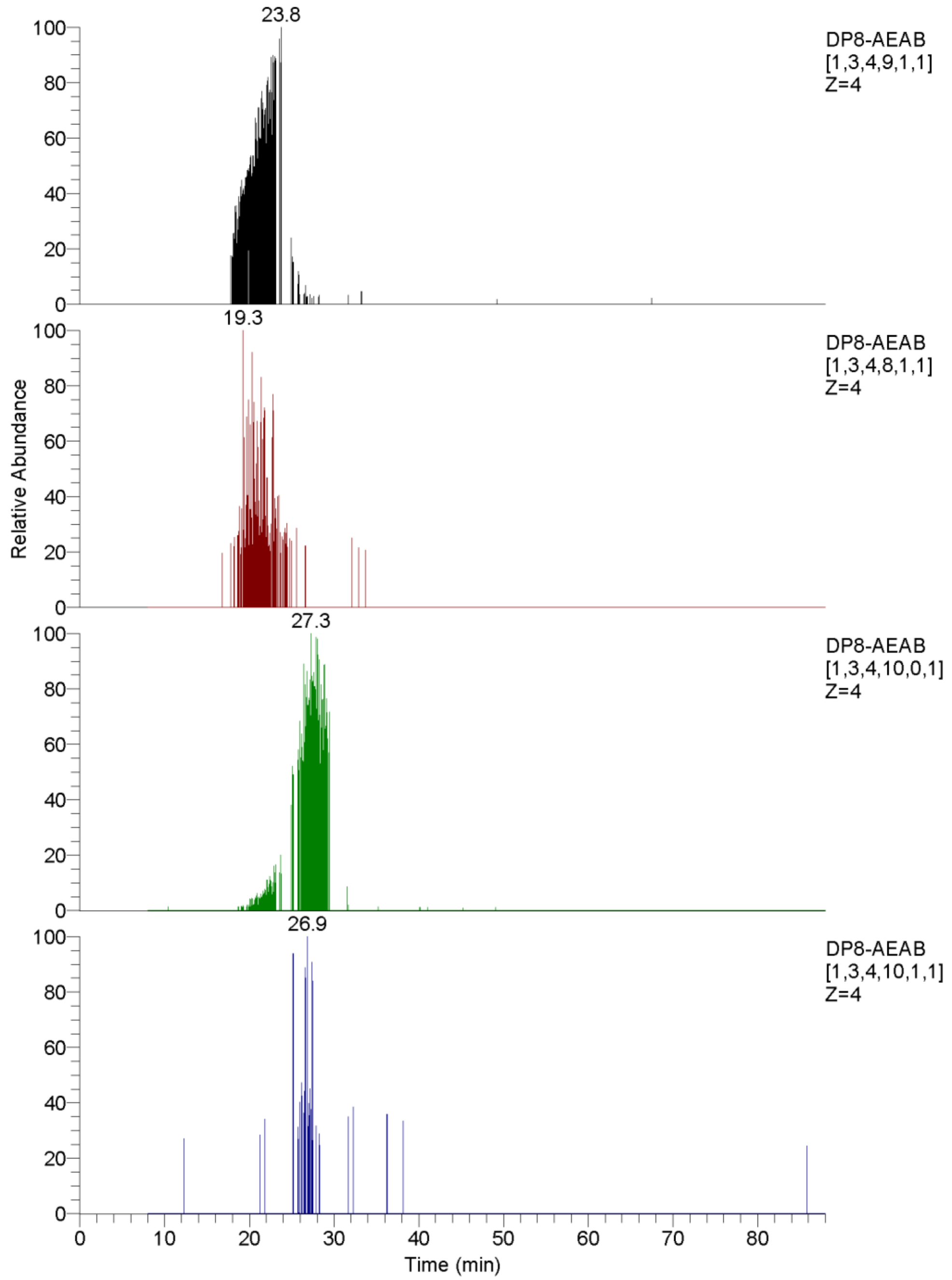


**Figure S9.**

The HILIC-MS EICs of the top four possible octasaccharide compositions of enoxaparin dp8 from the IPRP 40-41 min fraction.

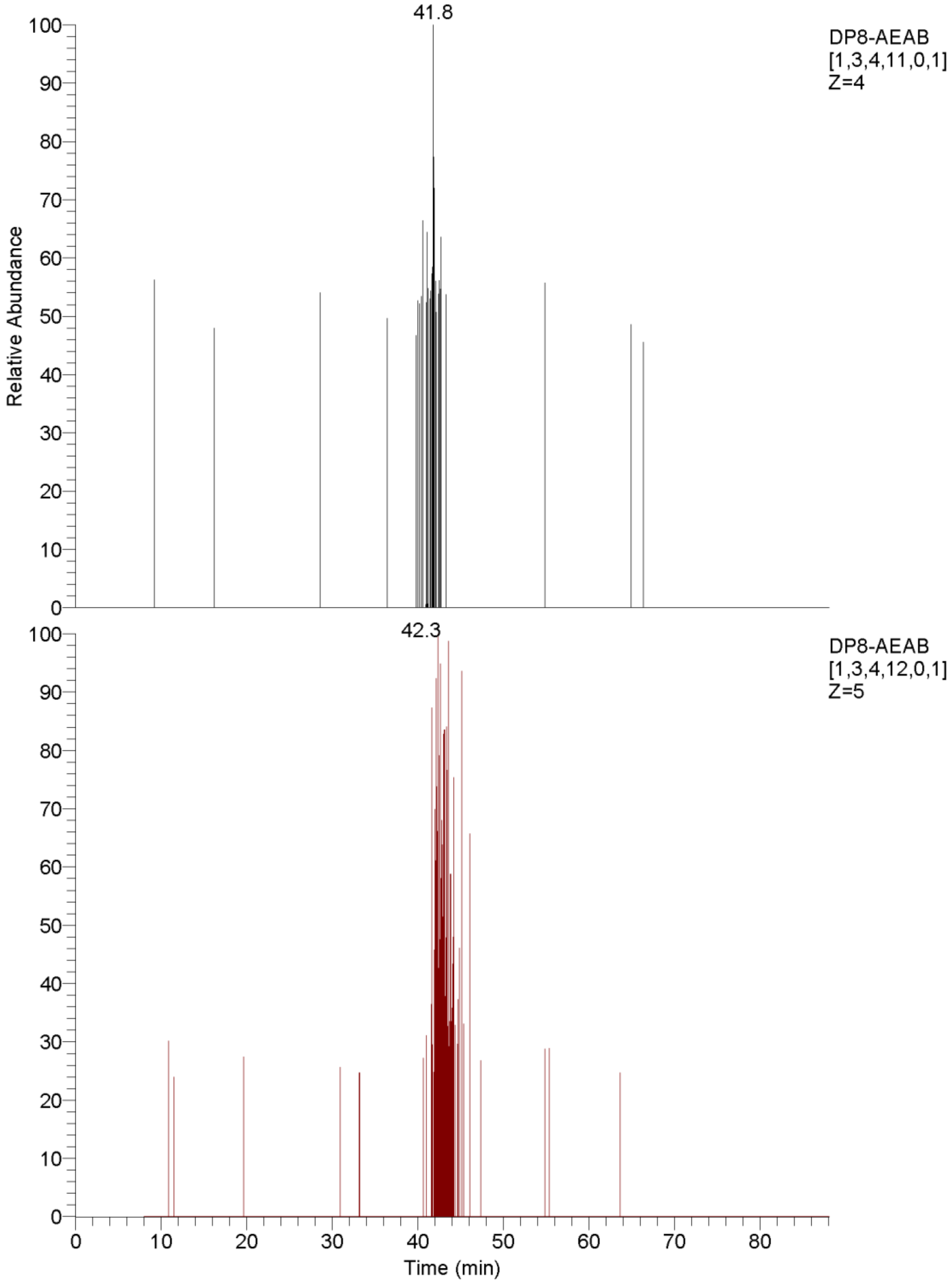


**Figure S10.**

The HILIC-MS EICs of the top four possible octasaccharide compositions of enoxaparin dp8 from the IPRP 58-61 min fraction.

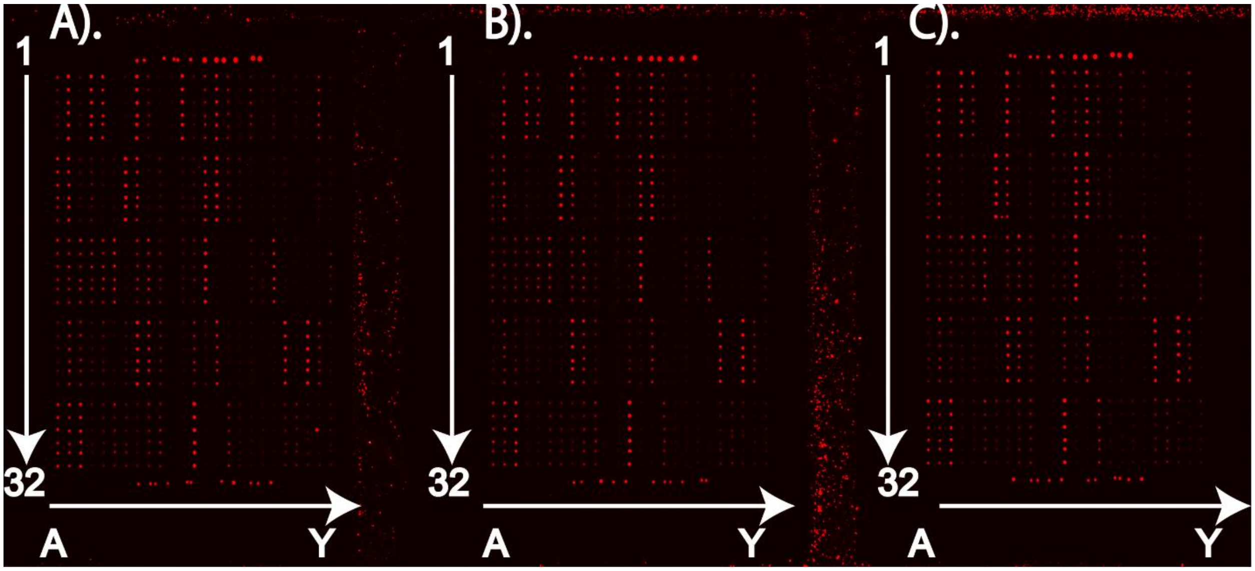


**Figure S11.**

The FGF2 binding affinity assay of 129 fractions of enoxaparin dp8. The blocks from left to right are binding affinities under various FGF2 concentrations, A). 1 µg/mL, B). 0.5 µg/mL, and C). 0.25 µg/mL, respectively. Fractions from No. 1 to No. 126 were eluents from the multi-dimensional separation method. Detail information of each fraction was listed in Table S1. Fractions No. 127 & 128 were unfractionated octasaccharide. Fraction No. 129 was synthetic octasaccharide (IdoA2S-GlcNS6S repeat) as a positive control.

| SAMPLE NO. | MICROARRAY POSITION | RT ON IPRP | RT ON HILIC | RFU(0.25 μg/mL) | RFU (0.5 μg/mL) | RFU (1 μg/mL) |
|---|---|---|---|---|---|---|
| **1** | Y2 to Y7 | 24-25.8 | 33-35 | 501.75±79.84 | 776.25±103.49 | 1138.75±262.32 |
| **2** | Y8 to Y13 | 24-25.8 | 36-38 | 1077.25±89.1 | 796.5±165.17 | 1333.5±129.61 |
| **3** | Y14 to Y19 | 26-26.8 | 32-35 | 4043.25±224.29 | 3152.75±410.21 | 3640.5±1046.13 |
| **4** | Y20 to Y25 | 26-26.8 | 36-37 | 1044.25±85.82 | 1026±117.7 | 1251.5±101.06 |
| **5** | Y26 to Y31 | 26-26.8 | 37-38 | 1987±122.36 | 1936.5±102.52 | 1985.5±136.37 |
| **6** | X2 to X7 | 27.5-28.3 | 32-35 | 5789±393.48 | 5348±391.62 | 7687.75±482.99 |
| **7** | X8 to X13 | 27.5-28.3 | 35-36 | 1801±167.97 | 1145.5±181.07 | 1826.75±216.73 |
| **8** | X14 to X19 | 27.5-28.3 | 36-38 | 506.75±137.2 | 526.75±89.53 | 671.5±164.94 |
| **9** | X20 to X25 | 27.5-28.3 | 38-40 | 5950.5±327.38 | 4417±137.75 | 5120±572.91 |
| **10** | X26 to X31 | 28.3-29.3 | 34-36 | 1343.75±163.91 | 1246.25±205.56 | 1471.75±479.78 |
| **11** | W2 to W7 | 28.3-29.3 | 36-37 | 830.5±153.33 | 876±242.39 | 1150.5±173.8 |
| **12** | W8 to W13 | 28.3-29.3 | 37-38 | 158±21.44 | 1±0 | 221.25±137.04 |
| **13** | W14 to W19 | 28.3-29.3 | 38-39 | 660.5±33.75 | 629±138.29 | 814.5±75.56 |
| **14** | W20 to W25 | 28.3-29.3 | 40-47 | 55108.25±1826.74 | 58939±5263.44 | 80225.5±5096.93 |
| **15** | W26 to W31 | 29.3-30.3 | 30-33 | 2680.25±138.6 | 2240.25±463.2 | 2052±323.35 |
| **16** | V2 to V7 | 29.3-30.3 | 35-36 | 1597.5±211.77 | 1294.25±171.94 | 2011.25±222.42 |
| **17** | V8 to V13 | 29.3-30.3 | 36-37 | 92.75±75.95 | 47.5±58.41 | 194.75±48.51 |
| **18** | V14 to V19 | 29.3-30.3 | 38-39 | 522±141.61 | 697±30.59 | 615.5±78.6 |
| **19** | V20 to V25 | 29.3-30.3 | 39-40 | 950.5±92.59 | 564.75±100.58 | 859±61.88 |
| **20** | V26 to V31 | 29.3-30.3 | 40-41 | 4276.5±254.65 | 3178±307.42 | 3660±141.82 |
| **21** | U2 to U7 | 30.3-30.7 | 36-37 | 234±115.65 | 321.25±96.62 | 371.25±111.17 |

| | | | | | | |
|---|---|---|---|---|---|---|
| **22** | U8 to U13 | 30.3-30.7 | 38-39 | 1342.75±193.94 | 1207.5±109.73 | 1392.75±178.17 |
| **23** | U14 to U19 | 30.3-30.7 | 39-40 | 1261.75±94.97 | 1232.5±64.49 | 1189.25±137.76 |
| **24** | U20 to U25 | 30.3-30.7 | 40-41 | 24018.5±734.28 | 17974.5±2251.11 | 27118.75±1382.64 |
| **25** | U26 to U31 | 30.7-31.6 | 30-35 | 3650±371.75 | 2402.25±281.94 | 3100.5±93.3 |
| **26** | T2 to T7 | 30.7-31.6 | 35-37 | 2852.25±56.7 | 2085.75±161.35 | 2893.5±56.98 |
| **27** | T8 to T13 | 30.7-31.6 | 37-40 | 1882.75±113.99 | 1243±20.77 | 1521±69.96 |
| **28** | T14 to T19 | 30.7-31.6 | 40-42 | 10622.25±394.52 | 8982.5±69.09 | 11092.25±991.56 |
| **29** | T20 to T25 | 31.6-32.3 | 37-39 | 1338.25±219.6 | 970.75±88.99 | 1223.75±102.86 |
| **30** | T26 to T31 | 31.6-32.3 | 39-40 | 1175.5±116.37 | 660.25±106.19 | 1058.25±83.83 |
| **31** | S2 to S7 | 32.3-32.6 | 35-37 | 1180.25±191.46 | 737±118.36 | 1271.75±172.03 |
| **32** | S8 to S13 | 32.3-32.6 | 37-38 | 891.5±133.15 | 669.75±116.61 | 1097.5±136.9 |
| **33** | S14 to S19 | 32.3-32.6 | 38-39 | 2152.25±139.31 | 1399.5±104.16 | 1952.75±150.46 |
| **34** | S20 to S25 | 32.3-32.6 | 39-40 | 591.5±117.74 | 440±88 | 570±132.18 |
| **35** | S26 to S31 | 32.6-33.7 | 34-36 | 1693.5±87.86 | 1158.75±60.74 | 1411.25±159.08 |
| **36** | R2 to R7 | 32.6-33.7 | 37-38 | 5248.75±178.13 | 4584.75±392.16 | 5414±307.31 |
| **37** | R8 to R13 | 32.6-33.7 | 38-39 | 1317.25±338.59 | 830.75±105.77 | 1293.25±226.78 |
| **38** | R14 to R19 | 32.6-33.7 | 40-41 | 3878.5±598.2 | 2586.75±204.71 | 2922.5±301.6 |
| **39** | R20 to R25 | 33.7-34.4 | 35-37 | 1400.25±60.11 | 1027.5±385.94 | 1342±104.9 |
| **40** | R26 to R31 | 33.7-34.4 | 37-39 | 1129.75±135.02 | 790.5±79.12 | 1066.25±116.97 |
| **41** | Q2 to Q7 | 33.7-34.4 | 39-40 | 957.25±116.06 | 905.75±127.84 | 1115.25±166.21 |
| **42** | Q8 to Q13 | 33.7-34.4 | 40-42 | 2780.5±216.3 | 2192.5±88.93 | 2839±128.11 |
| **43** | Q14 to Q19 | 34.4-35 | 36-38 | 470±140.12 | 626.25±84.16 | 615±30.7 |
| **44** | Q20 to Q25 | 34.4-35 | 38-40 | 1125±165.2 | 773.5±288.86 | 1030±155.94 |
| **45** | Q26 to Q31 | 34.4-35 | 40-42 | 1753.75±173.68 | 894.5±87.28 | 1407±65.49 |

| | | | | | | |
|---|---|---|---|---|---|---|
| **46** | P2 to P7 | 35.8-36.8 | 34-37 | 4714.25±319.29 | 4290.25±342.5 | 5162±321.72 |
| **47** | P8 to P13 | 35.8-36.8 | 37-39 | 3145±485.69 | 2613.75±265.93 | 4177±276.19 |
| **48** | P14 to P19 | 35.8-36.8 | 39-40 | 1111±277.14 | 703.5±50.18 | 812.25±101.75 |
| **49** | P20 to P25 | 35.8-36.8 | 40-41 | 1235.25±208.63 | 967.25±186.43 | 1281.5±101.1 |
| **50** | P26 to P31 | 35.8-36.8 | 41-42 | 6636±184.92 | 4520.25±368.44 | 5625±435.89 |
| **51** | O2 to O7 | 35.8-36.8 | 42-44 | 19208.5±1329.21 | 18374±1335.53 | 24615.75±2613.49 |
| **52** | O8 to O13 | 35.8-36.8 | 44-51 | 63592±1109.53 | 75754±1354.33 | 108554.5±3077.75 |
| **53** | O14 to O19 | 36.8-38 | 35-37 | 1320±107.01 | 847±121.95 | 1124.5±102.9 |
| **54** | O20 to O25 | 36.8-38 | 37-39 | 9550.75±302.8 | 6890±747.36 | 9883±757.83 |
| **55** | O26 to O31 | 36.8-38 | 39-40 | 846±84.15 | 647.5±102.05 | 763.75±106.46 |
| **56** | N2 to N7 | 36.8-38 | 40-42 | 5767.5±411.04 | 4578±340.16 | 6503±569.35 |
| **57** | N8 to N13 | 36.8-38 | 42-43 | 26893.75±1415.5 | 21920±1151.89 | 30451.25±1320.3 |
| **58** | N14 to N19 | 36.8-38 | 43-44 | 67833±2999.51 | 74839.25±2310.03 | 113813.75±2435.98 |
| **59** | N20 to N25 | 38-39 | 32-37 | 2591±262.86 | 1993.25±134.25 | 2302.25±499.88 |
| **60** | N26 to N31 | 38-39 | 38-39 | 1220.75±152.87 | 979.25±66.09 | 914±52.43 |
| **61** | M2 to M7 | 38-39 | 39-40 | 1861.25±232.46 | 1535±170.49 | 2239.5±321.7 |
| **62** | M8 to M13 | 38-39 | 40-41 | 2364.75±87.42 | 1933±152.7 | 2458.75±222.22 |
| **63** | M14 to M19 | 38-39 | 41-42 | 3332.75±163.07 | 2371.75±419.49 | 3643.25±368.54 |
| **64** | M20 to M25 | 38-39 | 42-43 | 5825.25±374.32 | 4048.5±413.23 | 5760±440.26 |
| **65** | M26 to M31 | 38-39 | 43-51 | 76164.5±3979.29 | 103855.25±3454.06 | 189050.75±4458.24 |
| **66** | L8 to L13 | 39-40.3 | 35-37 | 2766.25±114.81 | 2506.25±144.62 | 2388.5±258.62 |
| **67** | L14 to L19 | 39-40.3 | 37-38 | 2271±415.39 | 2043.25±135.57 | 2836.5±77.24 |
| **68** | L20 to L25 | 39-40.3 | 38-39 | 1194±249.75 | 1015.25±76.44 | 1012.25±175.68 |
| **69** | L26 to L31 | 39-40.3 | 39-40 | 1024.25±131.28 | 999.5±177.16 | 996±209.6 |

| | | | | | | |
|---|---|---|---|---|---|---|
| **70** | K2 to K7 | 39-40.3 | 41-43 | 1146.5±138.99 | 908.75±99.56 | 1248.5±150.57 |
| **71** | K8 to K13 | 41.5-41.9 | 37-40 | 285±25.49 | 384±26.07 | 283.5±40.02 |
| **72** | K14 to K19 | 41.5-41.9 | 41-42 | 371.25±104.01 | 239±145.23 | 328.75±128.98 |
| **73** | K20 to K25 | 41.5-41.9 | 42-43 | 182.25±9.93 | 32±33.79 | 220.25±77.41 |
| **74** | K26 to K31 | 41.5-41.9 | 43-44 | 199±91.01 | 156.5±102.08 | 297.5±128.49 |
| **75** | J2 to J7 | 42.5-44.3 | 40.5-41.5 | 783±94.32 | 483±142.09 | 902.5±186.39 |
| **76** | J8 to J13 | 42.5-44.3 | 41.5-42.5 | 3500.5±628.7 | 3112±153.68 | 3317.25±252.28 |
| **77** | J14 to J19 | 42.5-44.3 | 42.5-45.5 | 4754.25±501.33 | 3170±283.82 | 4657.75±667.9 |
| **78** | J20 to J25 | 44.3-45.1 | 35-39 | 4406.5±254.13 | 2999.25±396.25 | 4335.5±208.65 |
| **79** | J26 to J31 | 44.3-45.1 | 39.5-40.5 | 5034.5±562.32 | 4917.5±473.22 | 5719.75±856.29 |
| **80** | I2 to I7 | 44.3-45.1 | 41-42.5 | 2927.75±381.28 | 3683±253.95 | 4085±415.76 |
| **81** | I8 to I13 | 44.3-45.1 | 43-45.5 | 1441±42.44 | 1573.5±112.33 | 1942.75±191.08 |
| **82** | I14 to I19 | 46-46.8 | 36-38.5 | 7878±1242.91 | 7530.5±295.44 | 8228.25±1027.11 |
| **83** | I20 to I25 | 46-46.8 | 39 | 10584.5±1382.36 | 7914.25±304.28 | 11685.25±1196.76 |
| **84** | I26 to I31 | 46-46.8 | 40-41 | 3817.5±319.48 | 4532.5±300.6 | 4900.25±180.76 |
| **85** | H2 to H7 | 46-46.8 | 44-45 | 67618.25±1543.91 | 110939.75±3486.79 | 175298±4704.82 |
| **86** | H8 to H13 | 47-47.8 | 36-39 | 12239.5±1826.6 | 10766.5±1765.63 | 12541.75±2387.16 |
| **87** | H14 to H19 | 47-47.8 | 40-42 | 4857.75±827.27 | 5867.5±516.16 | 6782.75±499.43 |
| **88** | H20 to H25 | 47-47.8 | 42-44 | 38750±814.53 | 34947.75±4036.32 | 49977.75±5206.89 |
| **89** | H26 to H31 | 47.8-48.5 | 40-42 | 1836.25±230.71 | 2373±47.4 | 2666±323.57 |
| **90** | G2 to G7 | 47.8-48.5 | 42-44 | 1019±209.81 | 911±116.32 | 1368.25±132.02 |
| **91** | G8 to G13 | 47.8-48.5 | 44-51 | 70595±1402.31 | 97464.25±3515.06 | 126494.75±3986.91 |
| **92** | G14 to G19 | 49.3-49.7 | 35-40 | 914.75±230.88 | 749.5±206.63 | 1035.75±164.04 |
| **93** | G20 to G25 | 49.3-49.7 | 41-43 | 2038.25±269.94 | 1698±188.52 | 2881.75±261.04 |

| | | | | | | |
|---|---|---|---|---|---|---|
| **94** | G26 to G31 | 49.3-49.7 | 43 | 2544.75±134.93 | 3211±261.47 | 3896.5±243.29 |
| **95** | F2 to F7 | 49.3-49.7 | 44 | 289±82.98 | 320.5±71.39 | 416.5±58.9 |
| **96** | F8 to F13 | 50-50.7 | 36-39 | 930.75±137.66 | 913.5±163.34 | 1426.25±75.03 |
| **97** | F14 to F19 | 50-50.7 | 39-41 | 9532±591.61 | 9608.75±1315.25 | 12557.75±1293.63 |
| **98** | F20 to F25 | 50-50.7 | 42-45 | 1183.75±23.99 | 1048±48.14 | 1417.5±206.53 |
| **99** | F26 to F31 | 51-51.5 | 36-39 | 4297±205.58 | 3578.5±230.65 | 4834±730.22 |
| **100** | E2 to E7 | 51-51.5 | 40-42 | 8678.25±1992.54 | 8218.75±1164.3 | 14883.25±2357.93 |
| **101** | E8 to E13 | 51-51.5 | 43-44 | 823.75±129.52 | 1153±89.45 | 1550.75±315.92 |
| **102** | E14 to E19 | 51.5-52.6 | 36-40 | 5172.75±368.55 | 6535.25±413.96 | 7824±1176.67 |
| **103** | E20 to E25 | 51.5-52.6 | 40-41 | 2375.5±167.91 | 2217.5±486.69 | 3360.5±285.28 |
| **104** | E26 to E31 | 51.5-52.6 | 41-44 | 1789.5±208.54 | 1670.75±109.89 | 2994.25±267.12 |
| **105** | D2 to D7 | 51.5-52.6 | 45-47 | 24818.75±949.17 | 16694±1367.84 | 36302.75±5422.45 |
| **106** | D8 to D13 | 52.6-54.1 | 35-40 | 1745.25±308.91 | 2274±230.62 | 3807.75±331.58 |
| **107** | D14 to D19 | 52.6-54.1 | 42-44 | 4449±632.48 | 4719±343.93 | 7122.75±966.21 |
| **108** | D20 to D25 | 54.1-56.2 | 42-47 | 1988.5±308.55 | 2105±242.17 | 2621.75±264.86 |
| **109** | D26 to D31 | 56.8-57.5 | 40-44 | 1583.25±291.02 | 1693.25±181.59 | 1867.75±247.19 |
| **110** | C2 to C7 | 57.5-58.2 | 41-43 | 1239±395.83 | 1106.5±280.69 | 2270.5±233.56 |
| **111** | C8 to C13 | 58.2-60.9 | 43-46 | 1269.5±200.34 | 1603.75±134.59 | 2402±363.18 |
| **112** | C14 to C19 | 60.9-63.2 | 35-40 | 4211.5±604.74 | 4832.5±580.07 | 7703±522.63 |
| **113** | C20 to C25 | 60.9-63.2 | 40-42 | 4295.25±938.72 | 3744.25±292.42 | 6168.75±279.86 |
| **114** | C26 to C31 | 60.9-63.2 | 42-44 | 21949±1432.59 | 20198±1742.99 | 27313±2067.03 |
| **115** | B2 to B7 | 60.9-63.2 | 44-51 | 30974.5±1230.13 | 19824.25±9198.56 | 58380.5±4995.17 |
| **116** | B8 to B13 | 63.2-65 | 39-42 | 9943±1013.42 | 10512.75±687 | 21040±1110.53 |
| **117** | B14 to B19 | 63.2-65 | 42-45 | 3265.75±307.82 | 4487.5±330.38 | 7343±689.29 |

| | | | | | | |
|---|---|---|---|---|---|---|
| **118** | B20 to B25 | 63.2-65 | 45-48 | 6509±736.93 | 6282.75±869.74 | 8783.5±758.5 |
| **119** | B26 to B31 | 65-66.7 | 45-47 | 7658.5±980.24 | 6797±748.57 | 9483.5±1517.24 |
| **120** | A2 to A7 | 66.7-67.8 | 40-42 | 2527±616.27 | 1608.75±264.26 | 3894.25±340.78 |
| **121** | A8 to A13 | 66.7-67.8 | 42-44 | 3976.75±320.45 | 5380.25±197.94 | 9836.75±1807.99 |
| **122** | A14 to A19 | 67.8-69.5 | 40-45 | 4427±682.01 | 5049.75±644.14 | 7506.5±384.54 |
| **123** | A20 to A25 | 70-71.5 | 41-44 | 3053±560.3 | 2191.5±275.67 | 3332±385.38 |
| **124** | A26 to A31 | 72.2-74 | 45-50 | 10129.75±1098.31 | 7338.5±738.87 | 11186±258.92 |
| **125** | H32 to M32 | 74-75.5 | 48-54 | 14524.25±2727.49 | 12849.25±2733.86 | 10952.5±1548.79 |
| **126** | O32 to T33 | 75.5-77.5 | 46-54 | 16358.5±5386.22 | 11846±1456.25 | 13036.75±2580.73 |
| **127** | L2 to L7 | DP8-AEAB | | 54427.25±2242.52 | 63502.75±3128.2 | 95756.25±971.89 |
| **128** | H1 to M1 | DP8-AEAB | | 41424.5±1980.36 | 61537.5±1291.98 | 74164±9426.6 |
| **129** | N1 to S1 | Synthetic positive control | | 111723.75±13498.2 | 197024±6838.46 | 468389.75±8598.04 |

**Table S1.**

Enoxaparin fraction-FGF2 microarray results
